# Small fibre neuropathy in Fabry disease: a human-derived neuronal *in vitro* disease model

**DOI:** 10.1101/2023.08.09.552621

**Authors:** Thomas Klein, Julia Grüner, Maximilian Breyer, Jan Schlegel, Nicole Michelle Schottmann, Lukas Hofmann, Kevin Gauss, Rebecca Mease, Christoph Erbacher, Laura Finke, Alexandra Klein, Katharina Klug, Franziska Karl-Schöller, Bettina Vignolo, Sebastian Reinhard, Tamara Schneider, Katharina Günther, Julian Fink, Jan Dudek, Christoph Maack, Eva Klopocki, Jürgen Seibel, Frank Edenhofer, Erhard Wischmeyer, Markus Sauer, Nurcan Üçeyler

**Author notes:** Equal contribution (Thomas Klein, Julia Grüner). **Current affiliations:** Department of Women’s and Children’s Health, Karolinska Institutet, 17177 Stockholm, Sweden (Jan Schlegel). Institute for Biology and Environmental Science, University of Oldenburg, 26129 Oldenburg, Germany (Tamara Schneider). Bielefeld University, Medical Faculty Ostwestfalen-Lippe, Cellular Neurophysiology, 33615 Bielefeld, Germany (Erhard Wischmeyer). Institute of Molecular Biology & CMBI, University of Innsbruck, 6020 Innsbruck, Austria. **Correspondence to:** Prof. Nurcan Üçeyler, MD. Department of Neurology, University of Würzburg, Josef-Schneider-Str. 11, 97080 Würzburg, Germany. **Abbreviations:** 2i = 2 inhibitor cocktail; 5i = 5 inhibitor cocktail; AGAL = Agalsidase-beta; AP = Action Potential; BDNF = Brain-derived neurotrophic factor; BRN3A = Brain-specific homeobox/POU domain protein 3A; Ctrl = Control; DAPI = 4’,6-Diamidino-2-phenylindole; DMEM = Dulbecco’s modified Eagle medium; DMSO = Dimethyl sulfoxide; DRG = Dorsal root ganglia; EDTA = Ethylenediaminetetraacetic acid; EGTA = Egtazic acid; ERT = Enzyme replacement therapy; ERT = Enzyme replacement therapy; ExM = Expansion microscopy; FACS = Fluorescence activated cell sorting; FCS = Fetal calf serum; FD = Fabry disease; FdU = Floxuridine; FOXA2 = Forkhead box protein A2; FPQ = Würzburg Fabry Pain Questionnaire; Gb3 = Globotriaosylceramide; GFP = Green fluorescent protein; GDNF = Glial cell-derived neurotrophic factor; GLA = Alpha-galactosidase A; ICC = Immunocytochemistry; IENFD = Intraepidermal nerve fibre density; iPSC = Induced pluripotent stem cells; KO = Knockout; KSR = KnockOut medium; K_v_ = Voltage-gated potassium channel; Na_v_ = Voltage-gated sodium channel; NGFb = Nerve growth factor, beta subunit; OCT4 = Octamer-binding transcription factor 4; PAX6 = Paired box 6; PCR = Polymerase chain reaction; PFA = Paraformaldehyde; PNS = Peripheral nervous system; PRPH = Peripherin; qPCR = quantitative real-time PCR; QST = Quantitative sensory testing; RT = Room temperature; SCNxA = Sodium voltage-gated channel alpha subunit X; SIM = Structured illumination microscopy; SM22A = Smooth muscle protein 22-alpha; SOX2 = SRY-box transcription factor 2; SSEA4 = Stage-specific embryonic antigen-4; STxB = Shiga toxin 1, subunit B; TRPV1 = Transient receptor potential vanilloid type 1; TUJ1 = βIII-tubulin; XCI = X-chromosomal inactivation.

## Abstract

Acral burning pain triggered by fever, thermal hyposensitivity, and skin denervation are hallmarks of small fibre neuropathy in Fabry disease, a life-threatening X-linked lysosomal storage disorder. Variants in the gene encoding alpha-galactosidase A may lead to impaired enzyme activity with cellular accumulation of globotriaosylceramide (Gb3). To study the underlying pathomechanism of Fabry-associated small fibre neuropathy, we generated a neuronal *in vitro* disease model using patient-derived induced pluripotent stem cells from three Fabry patients and one healthy control. We further generated an isogenic control line via CRISPR/Cas9 gene editing. We subjected iPSC to targeted peripheral neuronal differentiation and observed intra-lysosomal Gb3 accumulations in somas and neurites of Fabry sensory neurons using super-resolution microscopy. At functional level, patch-clamp analysis revealed a hyperpolarizing shift of voltage-gated sodium channel steady-state inactivation kinetics in Fabry cell lines as compared to the healthy control. Moreover, we demonstrate a drastic increase in Fabry sensory neuron Ca^2+^ levels at 39°C mimicking clinical fever (p < 0.001). This pathophysiological phenotype was accompanied by thinning of neurite calibres in sensory neurons obtained from Fabry patients compared to healthy control cells (p < 0.001). Linear-Nonlinear cascade models fit to spiking responses revealed that Fabry cell lines exhibit altered single neuron encoding properties relative to control. We further observed jam of mitochondrial trafficking at sphingolipid accumulations within Fabry sensory neurites utilizing a click-chemistry approach together with mitochondrial dysmorphism compared to healthy control cells. We pioneer insights into the cellular mechanisms contributing to pain, thermal hyposensitivity, and denervation in Fabry small fibre neuropathy, and pave the way for further mechanistic *in vitro* studies in Fabry disease and the development of novel treatment approaches.

## Introduction

Fabry disease is an X-linked lysosomal storage disorder that is caused by variants in the gene encoding alpha-galactosidase A (*GLA*).^1^ Impairment of GLA activity results in cellular accumulation of sphingolipids, mainly globotriaosylceramide (Gb3).^2^ The clinical phenotype spans a spectrum from classic Fabry disease, which is an age-dependent multiorgan disorder starting in early childhood, to late-onset symptom manifestation in adulthood and with often milder symptoms. The main neurological manifestation of Fabry disease is small fibre neuropathy,^3^ which is characterised by episodic acral and triggerable burning pain, thermal hyposensitivity, and peripheral denervation both in men and women.^4^

The pathophysiology of small fibre neuropathy in Fabry disease is incompletely understood. Studies in Fabry animal models such as the *GLA* knockout (KO) mouse^5^ show high caspase activity in sensory neurons and altered ion channel function.^6^ Pain-related ion channels, such as members of the transient receptor potential vanilloid family and the family of voltage-gated sodium channels, are dysregulated.^7^ Further, gene expression related to lysosomes and ceramide metabolism is upregulated, whereas immune-related pathways are downregulated in *GLA* KO mice compared to wildtype.^8,9^ Although literature on the intracellular signalling pathways is sparse, a link between Ca^2+^ homeostasis and Fabry-associated symptoms has been proposed in murine sensory neurons^10^ and human cardiomyocytes derived from induced pluripotent stem cells (iPSC).^11^ Incubation of murine dorsal root ganglia (DRG) with lyso-Gb3, the deacylated form of Gb3, led to increased Ca^2+^ influx and iPSC-derived cardiomyocytes of patients with Fabry disease, increasing cytosolic Ca^2+^ transients. Also, the Ca^2+^-activated potassium channel KCa 3.1 was suggested to play a role in the pathogenesis of Fabry disease.^12^ However, there is hardly any data available on the mechanisms how functional alterations of cellular Gb3 may lead to Fabry pathophysiology in a human model. To comprehensively study these mechanisms in patients, human sensory neurons are needed which are not easily accessible *in vivo*.

Generation of sensory neurons from somatic cells via iPSC developed into a potent strategy to overcome this methodological roadblock.^13,14^ Multidimensional analysis of patient-derived sensory neurons provides in-depth insight into the molecular mechanisms underlying pain and sensory disturbance^15,16^ and paves the way for personalized treatment approaches.^17–19^ We pioneer a human *in vitro* model for Fabry disease and provide first evidence that GLA impairment is associated with altered neuronal properties as potential basis of small fibre neuropathy with pain, thermal hyposensitivity, and peripheral denervation. Our study is the first to link neuronal pathology with symptoms and signs of Fabry patients, opening avenues of unprecedented perspectives for future management of this life-threatening disease.

## Materials and methods

### Subjects

Our study was approved by the Würzburg Medical Faculty Ethics Committee (#135/15). Study participants gave written informed consent before inclusion and subjects’ consent was obtained according to the Declaration of Helsinki. We enrolled three patients (two men and one woman) with genetically approved Fabry disease in 2015 and 2016 via the Würzburg Fabry Center for Interdisciplinary Therapy (FAZIT), University of Würzburg. Additionally, we recruited a healthy adult male control subject.

### Clinical examination and pain assessment

Patients underwent complete neurological examination and were assessed with the Würzburg Fabry Pain Questionnaire (FPQ).^20^ Large fibre neuropathy was excluded by clinical examination and nerve conduction studies of the sural nerve following a standard procedure. Patients additionally underwent quantitative sensory testing (QST) for sensory profiles^21^ and skin punch biopsy.

### Skin punch biopsy and fibroblast cultivation

A 6-mm skin punch biopsy (Stiefel GmbH, Offenbach, Germany) was taken in local anesthesia from the lateral lower calf of all study participants.^22^ The skin sample was divided in two 3-mm halves. One half was used for immunohistochemistry and determination of the intraepidermal nerve fibre density (IENFD).^23,24^ From the second half, dermal fibroblasts were derived.^25^ Briefly, dermis and epidermis were mechanically separated and the dermal part was collected in fibroblast cultivation medium (DMEM/F12 + 100 U/ml penicillin 100 µg/ml streptomycin [pen/strep; both: Thermo Fisher Scientific, Waltham, MA, USA] + 10% fetal calf serum [FCS; Merck, Darmstadt, Germany]).

### Generation of iPSC

All cell lines were cultivated at 37°C with 5% CO_2_ (v/v). iPSC were generated using the StemRNA 3^rd^ Gen reprogramming kit (Reprocell, Maryland, USA) for all male cell lines (FD-1, FD-2, Ctrl) and the StemMACS mRNA reprogramming kit (Miltenyi Biotec, Bergisch Gladbach, Germany) for the female Fabry cell line (FD-3;^26^). Human dermal fibroblasts were seeded and transfected with the reprogramming cocktail for 4 (FD-1, FD-2, Ctrl) or 12 consecutive days (FD-3). Putative iPSC colonies were picked, expanded, and characterised (see below). Cells were cultivated on hESC-qualified Matrigel (Corning, Corning, USA) in StemMACS iPS-Brew XF cultivation medium (Miltenyi Biotec, Bergisch Gladbach, Germany) supplemented with 100 U/ml pen/strep (Thermo Fisher Scientific, Waltham, MA, USA). Cells were passaged twice a week using 2 mM ethylenediaminetetraacetic acid in phosphate buffer saline (EDTA/PBS; Thermo Fisher Scientific, Waltham, USA; Merck, Darmstadt, Germany) adding 10 µM Y27632 (Miltenyi Biotec, Bergisch Gladbach, Germany) for the first 24 hours after splitting and daily change of medium.

### iPSC characterisation

iPSC clones were extensively characterised.^27^ Putative iPSC were analyzed for the expression of the pluripotency-associated markers octamer-binding transcription factor 4 (OCT4, Santa Cruz Biotechnology, Dallas, USA), TRA-1-60 (Millipore, Burlington, MA, USA), and stage-specific embryonic antigen-4 (SSEA4, R&D Systems, Minneapolis, USA) using immunocytochemistry (ICC), with the latter two additionally analyzed by flow cytometry with suitable isotype antibodies (all: Miltenyi Biotec, Bergisch Gladbach, Germany) and unstained controls. To prove pluripotency, iPSC were differentiated into cells of all three germ layers using the StemMACS Trilineage Differentiation kit (Miltenyi Biotec, Bergisch Gladbach, Germany). Briefly, iPSC were seeded and cultivated with chemically-defined medium driving differentiation into each germ layer. Cells were analyzed via ICC for the expression of smooth muscle protein 22-alpha (SM22A, Abcam, Cambridge, UK), ectodermal paired box 6 and SRY-box transcription factor 2 (PAX6/SOX2, Biolegend, San Diego, CA, USA / R&D Systems, Minneapolis, MN, USA), and forkhead box protein A2 (FOXA2, Santa Cruz Biotechnology, Dallas, TX, USA) to verify mesodermal, ectodermal, and endodermal identity. To exclude chromosomal aberrations, karyotypes were evaluated using G-banding (FD-1, FD-2: Cell Guidance Systems, Cambridge, UK; FD-3: Creative Bioarray, Shirley, NY, USA; Ctrl: Institute for Human Genetics, University of Würzburg). Mutation analysis was done using PCR and Sanger sequencing (Eurofins Genomics, Ebersberg, Germany). GLA activity in iPSC was determined with the Alpha-Galactosidase Activity Assay Kit (Abcam, Cambridge, UK) at 42°C. Supernatant from iPSC was regularly screened for *Mycoplasma* DNA contamination via PCR.

### Isogenic Fabry cell line

We generated an isogenic Fabry cell line from the healthy control (ISO-FD) by CRISPR/Cas9 gene editing.^28^ A single-guide RNA (sgRNA) targeting *GLA* exon 7 was designed using the CHOPCHOP web tool^29^ and cloned into a plasmid carrying *S. pyogenes* Cas9 fused to 2A-GFP (pSpCas9(BB)-2A-GFP, Addgene, Watertown, MA, USA). The construct was transfected into iPSC using Lipofectamine Stem Transfection Reagent (Thermo Fisher Scientific, Waltham, MA, USA). GFP-positive cells were isolated via fluorescent activated cell sorting to obtain monoclonal lines and screened for successful gene editing by substrate staining with labelled Shiga toxin 1, subunit B (STxB), and Sanger sequencing. To verify enzyme dysfunction, GLA activity was measured. To ensure post-editing integrity of ISO-FD iPSC, basic characterisation comprising pluripotency marker expression, three-germ-layer differentiation, and karyotype analysis was repeated.

### Immunoreaction and expansion microscopy

Immunoreactions were performed following established protocols and depending on target location, cell type, and sample type. In brief, samples were fixed with 4% paraformaldehyde (PFA; Electron Microscopy Sciences, Hatfield, USA), blocked and permeabilized, if applicable, incubated with primary antibodies overnight, immunoreacted with matching secondary antibodies, and mounted for analysis. Fluorescent image acquisition and post processing followed determined rules (Supplementary Methods). Expansion microscopy was performed following published protocols (Supplementary Methods; Supplementary Fig. 1 and 2).^30,31^ For a list of antibodies, see Supplementary Table 1.

### Sensory neuron differentiation

Following a published protocol,^32^ iPSC on day −2 were dissociated and seeded into growth factor reduced Matrigel-(Corning, Corning, NY, USA) coated 6-well plates with a density of 120,000 cells/cm² and cultivated for 2 days adding 10 µM Y27632 for the first 24 h and daily medium changes. On day 0, medium was switched to KnockOut medium (KSR; KnockOut DMEM/F12 + 2 mM GlutaMAX + 15% KnockOut Serum Replacement + 100 µM 2-mercaptoethanol + 0.1 mM minimum essential medium non-essential amino acids + 100 U/ml Pen/Strep [all: Thermo Fisher Scientific, Waltham, MA, USA]), spiked with a two-inhibitor cocktail (2i) containing 100 nM LDN-193189 (Stemcell Technologies, Vancouver, Canada) and 10 µM SB-431542 (Miltenyi Biotec, Bergisch Gladbach, Germany). Starting from day 2, medium was supplemented with 2i and additionally a three inhibitor (3i) cocktail (10 µM SU-5402, 10 µM DAPT [both: Sigma Aldrich, St. Louis, MO, USA], and 3 µM CHIR-99021 [Axon Medchem, Groningen, Netherlands]). Starting at day 4, KSR medium was replaced by N2 medium (N2; DMEM/F12 GlutaMAX + 1X B-27 Plus Supplement + 1X N-2 Supplement + 100 U/ml pen/strep [all: Thermo Fisher Scientific, Waltham, MA, USA]), supplemented with 2i + 3i in 25% increments every two days until day 10. On day 10, cells were washed, detached with TrypLE Express (Thermo Fisher Scientific, Waltham, MA, USA), centrifuged and seeded in growth factor reduced Matrigel-coated 6-well plates, or 24-well plates with a 1:2, or 1:2.5 ratio, respectively, in neuronal maturation medium [N2 medium + 20 ng/ml BDNF + 20 ng/ml GDNF + 20 ng/ml NGFb (all: Peprotech, Rocky Hill, NJ, USA) + 200 ng/ml ascorbic acid (Sigma-Aldrich, St. Louis, MO, USA)], spiked with 10 µM floxuridine (FdU, Santa Cruz Biotechnology, Dallas, TX, USA) to reduce residual proliferative cells. Half of the medium was changed once a week, depending on consumption. Neurons were cultivated for ≥ 5 weeks before analysis (“mature neurons”). Where applicable, neurons were detached using TrypLE Express (Thermo Fisher Scientific, Waltham, MA, USA) and replated into the required culture vessel using conditioned and fresh media with a 1:1 ratio and addition of floxuridine. Time in culture between replating and functional analysis was 1 week.

### Sensory neuron treatment

Mature neurons were incubated with 1.32 µg/ml agalsidase-beta^33^ (AGAL; Sanofi Genzyme; Cambridge, MA, USA) for 24 h and were analyzed to assess Gb3 load. Neurons were immunoreacted with βIII-tubulin (TUJ1, Abcam, Cambridge, UK) and STxB::555. Coverslips were scanned with a DMi8 fluorescence microscope (Leica Microsystems, Wetzlar, Germany). TUJ1^+^ cells were counted using the cell counter plugin for ImageJ (U.S. National Institutes of Health, Bethesda, MD, USA)^34^ and analyzed for STxB^+^ profiles. Coverslips from ≥ 3 independent differentiations per condition were analyzed for each Fabry disease cell clone.

### Patch-clamp analysis and characterisation of single-neuron encoding

Whole-cell patch-clamp recordings were carried out on five-to eight-weeks old sensory neurons. All measurements were performed at RT and at 39°C since heat is a main trigger of pain in Fabry disease. Current-clamp recordings served to analyze action potential parameters such as threshold potential, amplitude, half-width, and firing frequency. To analyze voltage-gated sodium (Na_v_) channel and voltage-gated potassium (K_v_) channel characteristics, voltage-clamp recordings were performed. Current densities as well as activation and inactivation kinetics were calculated (Supplementary Methods). To assess stimulus-encoding characteristics of single neurons, Linear-Nonlinear (LN) and Generalized Linear point process (GLM) models were fit to action potential trains elicited by current-clamp stimulation with broad spectrum Gaussian noise.

### Gene expression analysis

Total RNA was extracted using miRNeasy mini kit (Qiagen, Hilden, Germany). 250 ng RNA was reverse transcribed with MultiScribe reverse transcriptase (Thermo Fisher Scientific, Waltham, MA, USA). Quantitative real time PCR (qPCR) was performed using gene specific TaqMan probes (all: Thermo Fisher Scientific, Waltham, MA, USA; see Supplementary Table 2) with 8.75 ng cDNA as template for targets and endogenous control, in a duplex PCR approach on a Quantstudio 3 qPCR machine (Thermo Fisher Scientific, Waltham, MA; USA). Data were analyzed using the ΔΔC_t_ method, normalizing expression of markers in Ctrl-iPSC to 1 and calculating the relative gene expression accordingly, using *GAPDH* as housekeeping gene.

Human Voltage-Gated Ion Channel array plates (Thermo Fisher Scientific, Waltham, MA, USA) were loaded with pooled cDNA from two clones with two differentiations (FD-1, FD-2, Ctrl) or one clone with two differentiations (ISO-FD). Each well contained 9.26 ng cDNA and the reactions were performed on a Quantstudio 3. Data were quantified via ExpressionSuite Software (Thermo Fisher Scientific, Waltham, MA; USA) using the combined expression values of endogenous controls *18S*, *GAPDH*, *GUSB*, and *HPRT1* for ΔC_t_ normalization. Auto-threshold was applied for all targets based on the combined runs of all arrays and a cut-off C_t_ ≥ 33 was used to determine absence of a respective transcript. Expression is either depicted as ΔC_t_ or normalized to Ctrl neurons (relative expression, log2fold change). Principle component analysis was carried out via Clustvis^35^ with relative expression as input at default parameters.

### X-chromosome inactivation analysis and *GLA* transcription

X-chromosome inactivation (XCI) was analyzed using HhaI digestion, subsequent amplification, and fragment analysis (Supplementary Methods).

### Ca^2+^ imaging

Mature neurons were loaded with 2 µM Fluo-8 AM (Abcam, Cambridge, UK) for 1 h at 37°C and washed with conditioned neuronal medium. Images were acquired using a confocal laser scanning microscope (LSM700, Zeiss, Oberkochen, Germany) under physiological conditions utilizing a live-cell acquisition chamber (Tokai Hit, Shizuoka, Japan) with a sampling rate of 0.25 Hz for 8 min. For automated and objective analysis of neuronal activity over time, the image processing software Line Profiler (https://line-profiler.readthedocs.io/en/latest/) was used. Line Profiler applies a skeletonized algorithm^36^ to reduce expanded structures to one pixel width. The remaining pixel coordinates are fitted with a c-spline. This gives an analytical description of the structurès orientation and a suitable approximation for its centre. Line profiles are constructed perpendicularly to the derivative of the c-spline. The average of all line profiles is fitted with a Gaussian function

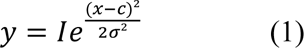

where c denotes the centre of the peak. The standard deviation *σ* and intensity *I* are saved in a text file for further evaluation. This process is repeated for each timeframe in all regions of interest and allows conclusions about the difference in activity between Fabry and Ctrl neurons.

### Mitochondrial mobility and morphology

To investigate mitochondrial mobility, mature Ctrl neurons were incubated with 1 µM of ω-N_3_ sphinganine (Cayman Chemical Company, Ann Arbor, MI, USA) overnight. The next day, cells were washed with maturation medium, and 1 µM BODIPY-PEG_4_-DBCO (Jena Bioscience, Jena, Germany) and 100 nM MitoTracker Deep Red (Thermo Fisher Scientific, Waltham, MA, USA) were added in maturation medium before incubation for 30 min in the incubator.^37^ Live-cell data were acquired using a Lattice-SIM microscope (Elyra 7, Zeiss, Oberkochen, Germany) with appropriate laser lines. As control conditions, neurons were only incubated with 1 µM BODIPY and 100 nM MitoTracker Deep Red (“Dye control”).

Mitochondrial morphology was analyzed after Tom20 antibody labeling. Photomicrographs were taken using a THUNDER Imager fluorescence microscope (Leica DMi8, Leica Microsystems, Wetzlar, Germany) and analyzed with ImageJ plugin Shape Descriptor.^38^ The morphology parameters “form factor” and “aspect ratio” were assessed.^39^ The form factor for branching was computed according to the formula

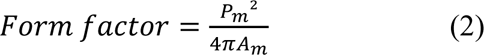

(P_m_ = outline length of mitochondrial area; A_m_ = mitochondrial area).

The aspect ratio was calculated as the ratio of the major and minor axis of an ellipse equaling the shape of the mitochondrion.

### Seahorse assay

Sensory neurons were seeded into Seahorse cell plates (Agilent, Santa Clara, CA, USA) and cultivated for six weeks. One hour before measuring, cell media were changed to Seahorse XF medium (Agilent, Santa Clara, CA, USA) supplemented with 5.5 mM glucose and 1 mM pyruvate. From this time point, cells were incubated without supply of 5% CO_2_. After measurement of baseline respiration sequential administration of 3 µM oligomycin (Sigma Aldrich, Saint Louis, MO, USA) was used to measure coupling efficiency, 1 µM FCCP (Sigma Aldrich, Saint Louis, MO, USA) for spare respiratory efficiency and 2 µM Rotenon (Sigma Aldrich, Saint Louis, MO, USA) + 1 µM antimycin (Sigma Aldrich, Saint Louis, MO, USA) for intoxication.

### Statistical analysis

Data analysis was performed with SPSS Statistics 27 (IBM, Armonk, NY, USA) with appropriate parametric or non-parametric tests, stated in the respective figure legends. Data were visualized using GraphPad PRISM Version 9.5 (GraphPad Software, Inc., La Jolla, CA, USA).

### Data availability

The data that support the findings of this study are available from the corresponding author, upon reasonable request.

## Results

### Clinical characterisation

We investigated two men (FD-1, FD-2) and one woman (FD-3) with genetically confirmed Fabry disease, and one man as Ctrl with wildtype *GLA*. Characteristics of the study participants is summarized in Supplementary Table 3. Both male patients carried pathogenic nonsense variants in *GLA* caused by a base substitution (FD-1) leading to a stop codon or a single-base deletion leading to a frame-shift (FD-2). FD-3 was diagnosed with a heterozygous missense variant located deeply within the GLA protein and classified as “buried mutation”.^40^ FD-1 showed a severe Fabry phenotype with cardiomyopathy, nephropathy, and Fabry-associated pain since early childhood. FD-1 also had further signs of Fabry-associated small fibre neuropathy, namely elevated thermal perception thresholds in QST and reduced IENFD of 4.9 fibres/mm (laboratory reference value: 9 ± 3 fibres/mm; Supplementary Fig. 3). In contrast, FD-2 did not have organ involvement and no signs or symptoms of small fibre neuropathy except for asymptomatic reduction of IENFD to 3.7 fibres/mm (Supplementary Fig. 3).^41^ FD-3 reported Fabry-associated pain attacks without further clinical signs of small fibre neuropathy. She had normal sensory perception thresholds in QST and normal skin innervation (8.5 fibres/mm). We further investigated Fabry and Ctrl skin cryosections and fibroblasts for Gb3 load using fluorescently labelled STxB.^42^ We found dense Gb3 deposits in the dermis and in fibroblasts of all three patients, which were absent in Ctrl (Supplementary Fig. 3).

During regular follow-up visits in 2021, symptoms and clinical signs had further deteriorated in FD-1 (i.e. six years after study inclusion, under AGAL treatment). In FD-2 (i.e. five years after study inclusion, without treatment), clinical status was unchanged to baseline visit. FD-3 was lost to follow-up.

### Fibroblasts from Fabry patients can be reprogrammed to iPSC retaining genotype and cellular phenotype

FD-1, FD-2, and ISO-FD iPSC lines expressed the pluripotency-associated proteins OCT4, TRA-1-60, and SSEA4 (Supplementary Fig. 4A). Directed differentiation of all lines into the three germ layers resulted in cells expressing SM22A, FOXA2, or PAX6, and SOX2 (Supplementary Fig. 4B). TRA-1-60 and SSEA4 expression was additionally analysed by flow cytometry with >75% of all TRA-1-60^+^/SSEA4^+^ cells (Supplementary Fig. 4C). G-banding showed a normal male karyotype (FD-1, FD-2, Ctrl, ISO-FD; 46,XY) after reprogramming and gene editing, respectively (Supplementary Fig. 4D). Patient-specific variants in *GLA* were confirmed for FD-1-iPSC (c.1069C>T, p.Q357X; hemizygous) and FD-2-iPSC (c.568delG; p.A190Pfs*1; hemizygous), whereas no variants were found in *GLA* of Ctrl-iPSC (Supplementary Fig. 4E). In ISO-FD-iPSC, the *GLA* variant c.1091_1093del (p.S364del, hemizygous) was generated by CRISPR/Cas9 gene editing. Additionally, we employed FD-3 iPSC that were characterised as reported previously.^43^

### Fabry iPSC show persisting Gb3 accumulation only in cells of men, while iPSC of women undergo *in vitro* restitution by skewed XCI

While Ctrl-iPSC showed no Gb3 depositions (Fig. 1A), FD-1-, FD-2-, and ISO-FD-iPSC had numerous Gb3 accumulations (Fig. 1B - D). Surprisingly, female FD-3 iPSC displayed Gb3 depositions only in the early (Fig. 1E), but not late phase of cultivation (> passage 10; Fig. 1F). We hence analysed cells for XCI patterns as potential source for the loss of Gb3 during long-term cultivation. *GLA* cDNA sequencing showed the presence of only the wildtype (wt) allele on a transcriptional level in contrast to the genomic level, where both the wt and c.708G>C variant were detected (Fig. 1G). Methylation analysis of the androgen receptor gene (*AR*) revealed a skewed XCI pattern of 0:100, indicating a nonrandom XCI of one of the parental X-chromosomes rather than a random XCI, where the expected XCI ratio usually follows a 70:30 to 50:50 distribution (Fig. 1H). Since no affected male relative carrying the *GLA* missense mutation was available for segregation analysis, the polymorphic (CAG)_n_ repeat sequence of the fully inactivated *AR* allele could not be assigned to the X-chromosome carrying the mutated *GLA* allele nor to the wt allele. However, the lack of Gb3 accumulations, the loss of mRNA expression from the mutated allele, and the skewed XCI pattern together highly suggest a selective and 100% inactivation of the X-chromosome carrying the *GLA* missense variant. Therefore, FD-3 line was excluded from further experiments.

**Figure 1:**
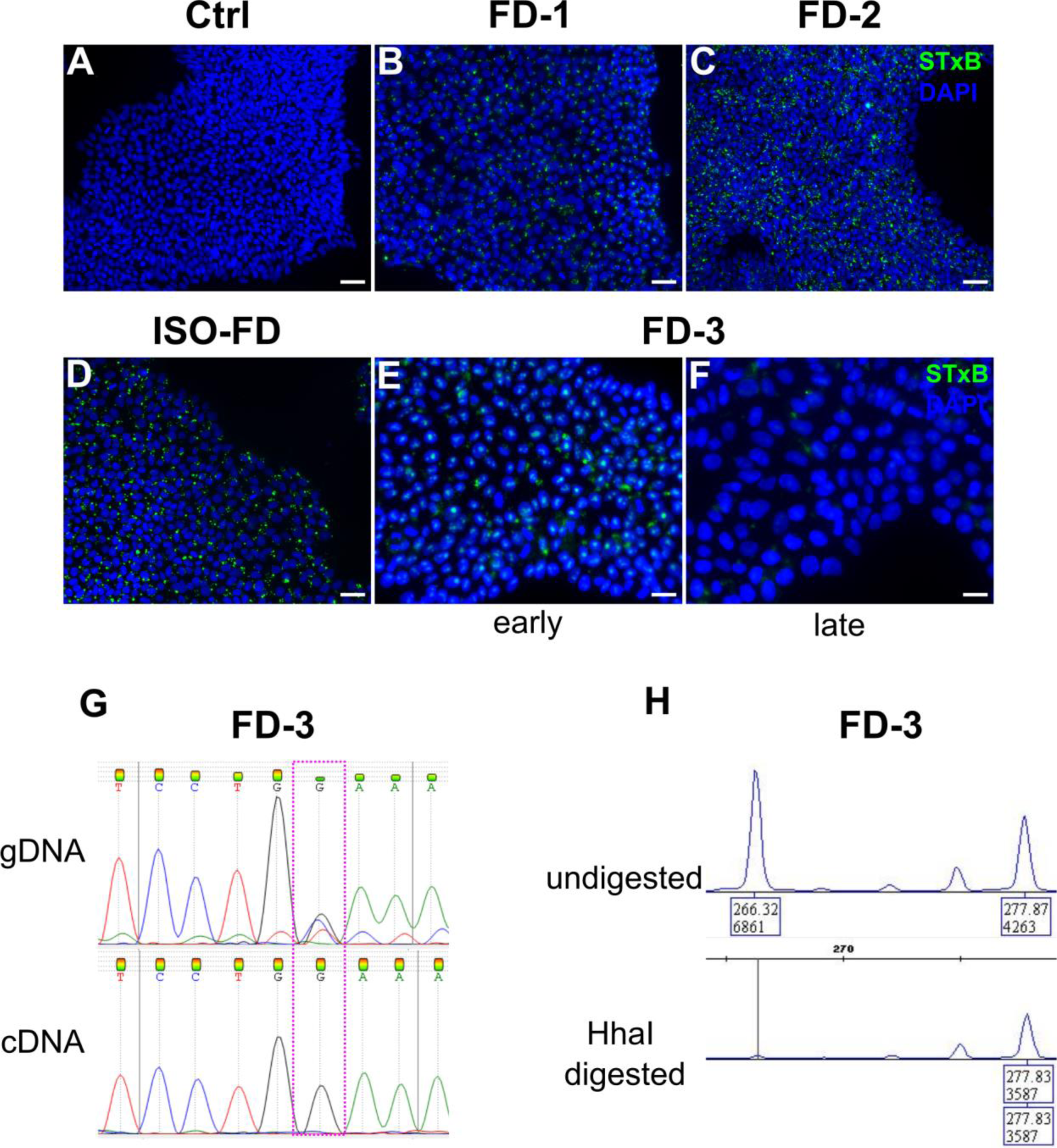
Gb3 accumulations in iPSC and skewed X-chromosome inactivation. (**A**) Ctrl-iPSC show no Gb3 accumulations. Scale bar: 50 µm. (**B**) - (**D**) FD-1, FD-2, and ISO-FD iPSC display numerous Gb3 accumulations. Scale bars: 50 µm. (**E**) iPSC from female FD-3 lines show Gb3 accumulations in early passages. Scale bar: 25 µm. (**F**) Gb3 accumulations are lost in iPSC of FD-3 during cultivation (> 10 passages). Scale bar: 25 µm. (**G**) Analysis of gDNA via Sanger sequencing shows the expected heterozygous disease-associated mutation, whereas analysis of cDNA from late FD-3 cells (> 10 passages) shows the wildtype sequence. (**H**) Analysis of X-chromosomal inactivation in late-passage (> 10 passages) by methylation-sensitive fragment analysis of the polymorphic (CAG)_n_ repeat in the *AR* gene. FD-3 cells show a complete shift towards one allele. **Abbreviations:** Ctrl = control, DAPI = 4’,6-diamidino-2-phenylindole, FD-1, FD-2, FD-3 = patients with Fabry disease, iPSC = induced pluripotent stem cells, STxB = shiga toxin 1, subunit B.

Mimicking the clinical phenotype, GLA enzyme activity was virtually absent in FD-1-iPSC (10.78 nmol/h/mg) and below the assay’s detection threshold for FD-2 (< 0.006 nmol/h/mg; Supplementary Fig. 4F) compared to normal activity in Ctrl-iPSC (508.98 nmol/h/mg; Supplementary Fig. 4F). GLA activity was also absent in ISO-FD-iPSC (14.57 nmol/h/mg; Supplementary Fig. 4F) confirming the disrupting effect of CRISPR/Cas9 gene editing on enzyme function.

### Functional sensory neurons derived from Fabry iPSC exhibit lysosomal Gb3 deposits cleaved by AGAL

During differentiation, cellular morphology changed to neuron-like cells starting on day five. After ten days, a ganglion-like morphology was visible with also non-neuronal cells present (Fig. 2A). Upon splitting and treatment with FdU, the number of non-neuronal cells decreased substantially (Fig. 2B). Immunocytochemistry showed expression of TUJ1 (Fig. 2C), peripherin (PRPH; Fig. 2C), Na_v_ 1.8 (Fig. 2D), substance P (TAC1; Fig. 2E), and transient receptor potential vanilloid type 1 (TRPV1; Fig. 2F) in mature neurons. Higher expression of neuronal markers *TUJ1*, *PRPH*, *brain-specific homeobox/POU domain protein 3A (BRN3A), tropomyosin receptor kinase A (TRKA), islet-1 (ISL1), TRPV1, TAC1*, and *Na_v_ 1.7* was confirmed in differentiated cells compared to iPSC via qRT-PCR (Fig. 2G). Expression of Na_v_ 1.8 was detected in neurons (Ct < 35), but not in iPSC (Ct ≥ 35).

**Figure 2:**
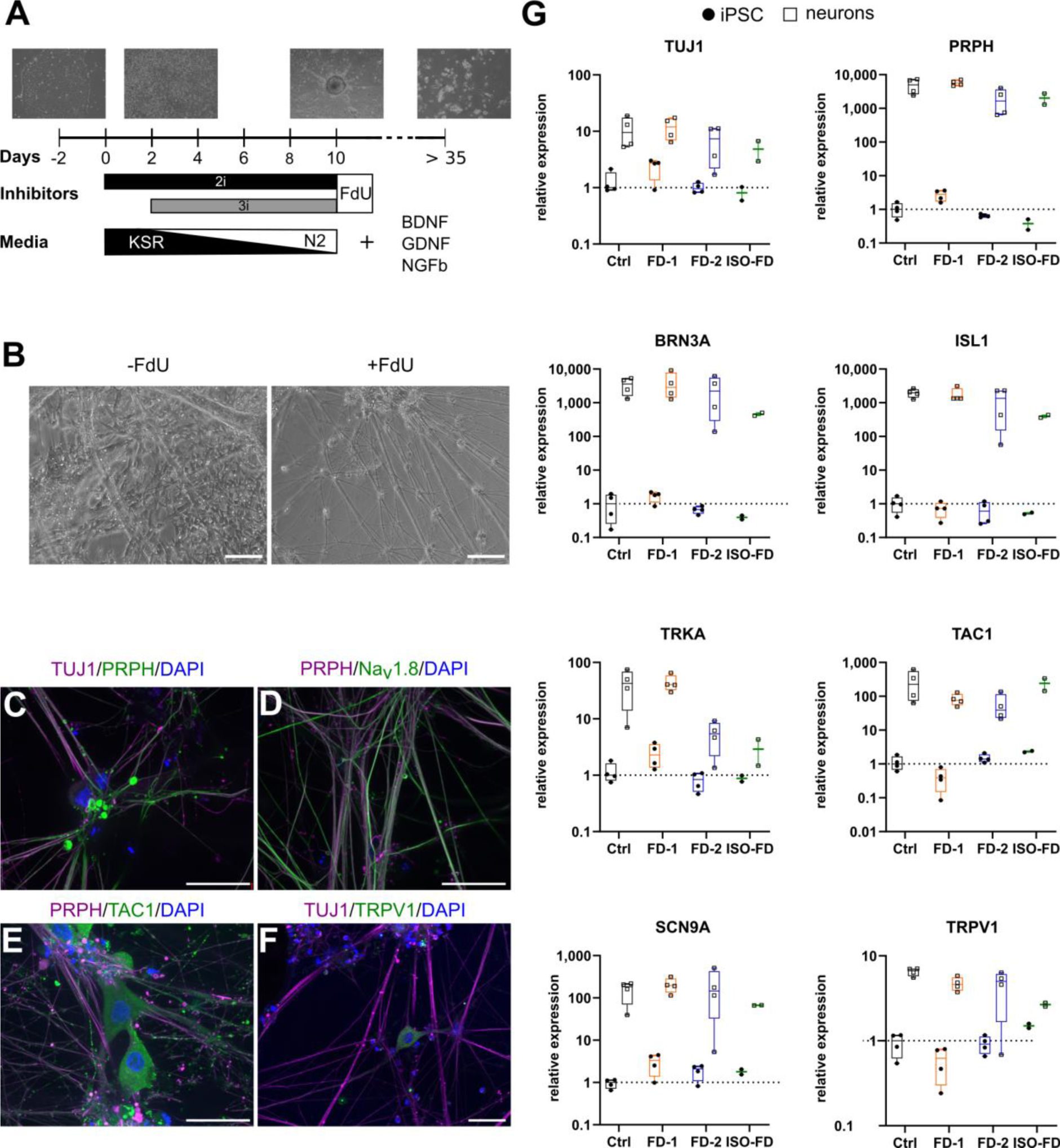
Differentiation strategy and characterisation of iPSC-derived sensory neurons. (**A**) Schematic overview of the differentiation from iPSC to sensory neurons depicting the time frame and media composition. (**B**) Incubation with FdU leads to a decrease in non-neuronal cell populations. Scale bars: 50 µm. (**C**) iPSC-derived sensory neurons show expression of neuronal (TUJ1) and peripheral (PRPH) markers. Scale bar: 25 µm. (**D**) Co-expression of TUJ1 and nociceptive Na_v_1.8. Scale bar: 25 µm. (**E**) Co-expression of TUJ1 and the peptidergic nerve fibre marker TAC1. Scale bar: 25 µm. (**F**) TRPV1, a marker for nociceptors, is co-expressed with TUJ1. Scale bar: 25 µm. (**G**) qPCR analysis of total mRNA from iPSC-derived neurons confirmed the expression of genes associated with the sensory and nociceptive lineage. Data are represented as mean ± SD. For (**G**), pooled data from n = 2 clones/line, obtained from two individual differentiations each. For ISO-FD, pooled data from n = 1 clone from n = 2 individual differentiations. **Abbreviations:** BDNF = brain-derived neurotrophic factor, FdU = Floxuridine, GDNF = glial cell-derived neurotrophic factor, NGFb = nerve growth factor, beta subunit, PRPH = peripherin, SCN9A = sodium voltage-gated channel alpha subunit 9, TAC1 = Tachykinin Precursor 1, TRPV1 = Transient receptor potential vanilloid type 1, TUJ1 = βIII-tubulin.

We characterised cellular Gb3 distribution in iPSC and sensory neurons by expansion microscopy and structured illumination microscopy co-applying antibodies against lysosomal-associated membrane protein 1 (LAMP1). In Ctrl iPSC, no Gb3 was detected (Fig. 3A), whereas in iPSC of FD-1 (Fig. 3B) and FD-2 (Fig. 3C), numerous intra-lysosomal Gb3 accumulations were present. Quantification of Gb3 accumulations in sensory neurons showed that Ctrl sensory neurons were free of Gb3 accumulations (Fig. 3D, E), while neurons of FD-1, FD2, and ISO-FD showed dense Gb3 depositions (Fig. 3F – H; 3I, J). Gb3 deposits were cleaved by 27% in FD-1 (p < 0.001), 22% in FD-2 (p < 0.01), and 16% in ISO-FD sensory neurons upon 24-h AGAL incubation (Fig. 3D).

**Figure 3:**
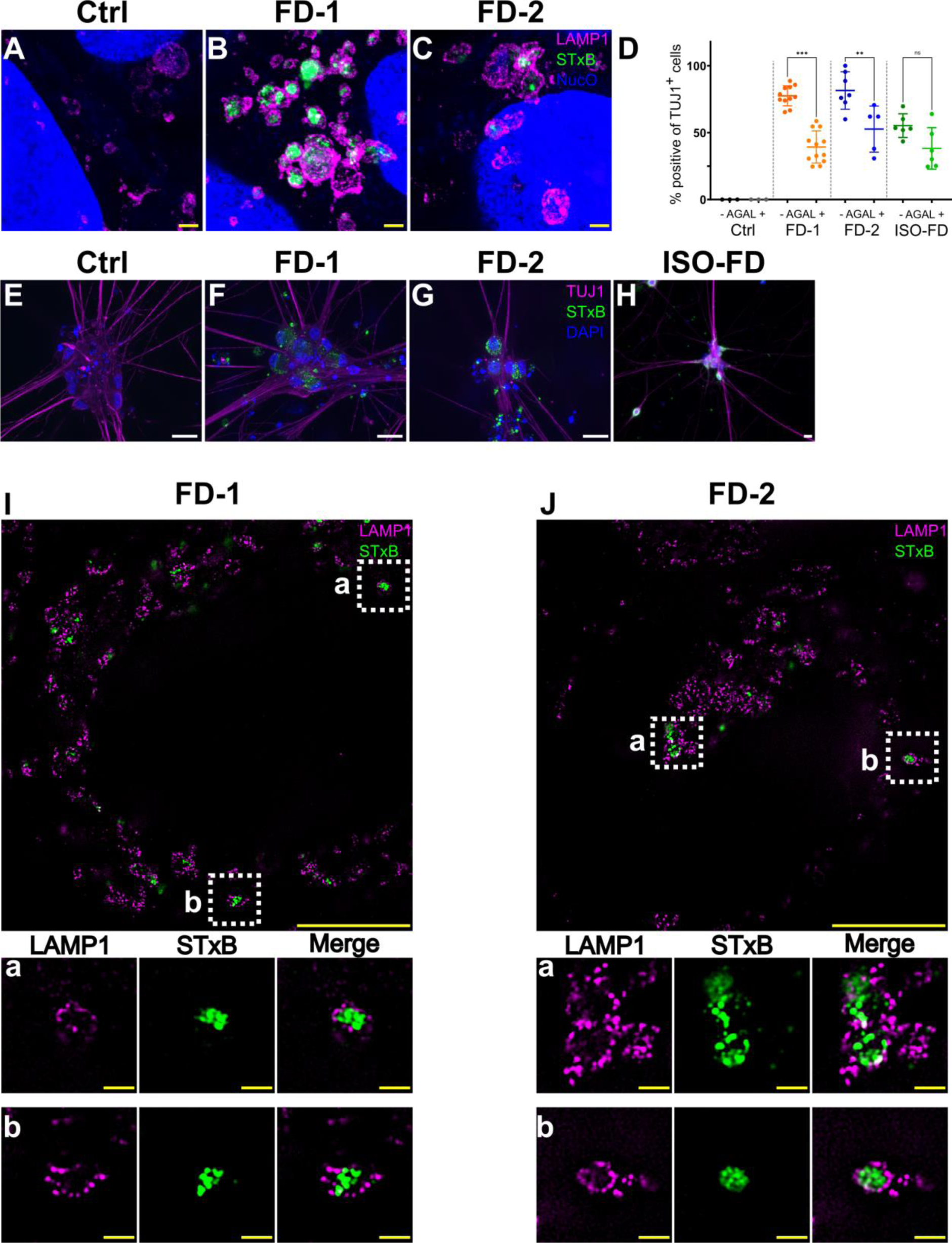
AGAL incubation and in-depth analysis of Gb3 accumulations. (**A**) Lysosomes of expanded Ctrl-iPSC show no accumulations. Scale bar: 1 µm (expansion factor corrected). (**B**) Lysosomes of expanded FD-1 iPSC show prominent intra-lysosomal Gb3 accumulations. Scale bar: 1 µm (expansion factor corrected). (**C**) Lysosomes of expanded FD-2 iPSC show prominent intra-lysosomal Gb3 accumulations. Scale bar: 1 µm (expansion factor corrected). Nuclear Orange (NucO) was used to visualize the nuclei. Acquired with LSM700 (confocal). (**D**) Incubation with AGAL reduced Gb3 in FD-1 (untreated: n = 2 clones; clone 1 = 457 cells, clone 2 = 1050 cells; +AGAL: n = 2 clones; clone 1 = 556 cells, clone 2 = 1030 cells) and FD-2 (untreated: n = 2 clones; clone 1 = 341 cells, clone 2 = 300 cells; +AGAL: n = 2 clones; clone 1 = 476 cells, clone 2 = 150 cells) neurons. AGAL incubation decreased Gb3 deposits also in ISO-FD, although statistics did not reach significance (untreated: n = 1 clone, n = 628 cells; +AGAL: n = 1 clone, n = 610 cells). Ctrl neurons did not show any accumulations before (n = 1 clone, 150 cells) and after (n = 1 clone, 150 cells) AGAL incubation. Data are represented as mean ± SD. One-way ANOVA followed by Sidak’s multiple comparison correction. (**E**) No Gb3 depositions were detected in Ctrl neurons. Scale bar: 25 µm. (**F**) - (**H)** FD-1, FD-2, and ISO-FD-neurons showed massive Gb3 accumulations. Scale bars: 25 µm. (**I**) Super-resolution image of an expanded FD-1 neuron. Scale bar: 5 µm (expansion factor corrected). (a), (b) ROIs cropped from (I) show intra-lysosomal Gb3 accumulations. Scale bars: 0.5 µm (expansion factor corrected). (**J**) Super-resolution image of an expanded FD-2 neuron. Scale bar: 5 µm (expansion factor corrected). (a), (b) ROIs cropped from (J) show intra-lysosomal Gb3 accumulations. Scale bars: 0.5 µm (expansion factor corrected). **Abbreviations:** AGAL = agalsidase-beta, Ctrl = control, DAPI = 4’,6-diamidino-2-phenylindole, FD-1, FD-2 = patients with Fabry disease, Gb3 = globotriaosylceramide, ISO-FD = isogenic Fabry line, ROI = region of interest, NucO = Nuclear Orange, PRPH = peripherin, STxB = shiga toxin 1, subunit B, TUJ1 = class III beta-tubulin. p < 0.05 = *; **p < 0.01 = **; p < 0.001 = ***. For expansion factors, see also Figures S2 and S3.

### Fabry sensory neurons show distinct voltage-gated ion channel expression profiles

IPSC-derived neurons differentiated towards the peripheral lineage were further characterised for mRNA expression patterns of pain-associated voltage-gated ion channels. Micro-array analysis demonstrated that Fabry sensory neuron expression patterns were overall distinct from Ctrl neurons, but also showed inter-individual diversity as illustrated by principle component analysis (Fig. 4A). The array revealed an independent signature for sensory neurons of FD-1 who also reported pain, while expression of FD-2 and ISO-FD neurons appeared clustered (Fig. 4A, B). Notably, among the 66/92 (72%) genes detected, *SCN9A* was expressed highest in all cell lines affirming their sensory nature (Fig. 4B). When analysing cell line signatures, all FD lines shared reduced expression of predominantly voltage-gated potassium channel family members (*KCND2, KCNJ11, KCNJ4, KCNK12*), yet FD-1 differed from both other FD lines and the control as indicated by increased mRNA expression of several channels (*KCNAB2, CACNG5, SCN3B, SCN7A, KCNB2*) (Fig. 4B, C).

**Figure 4:**
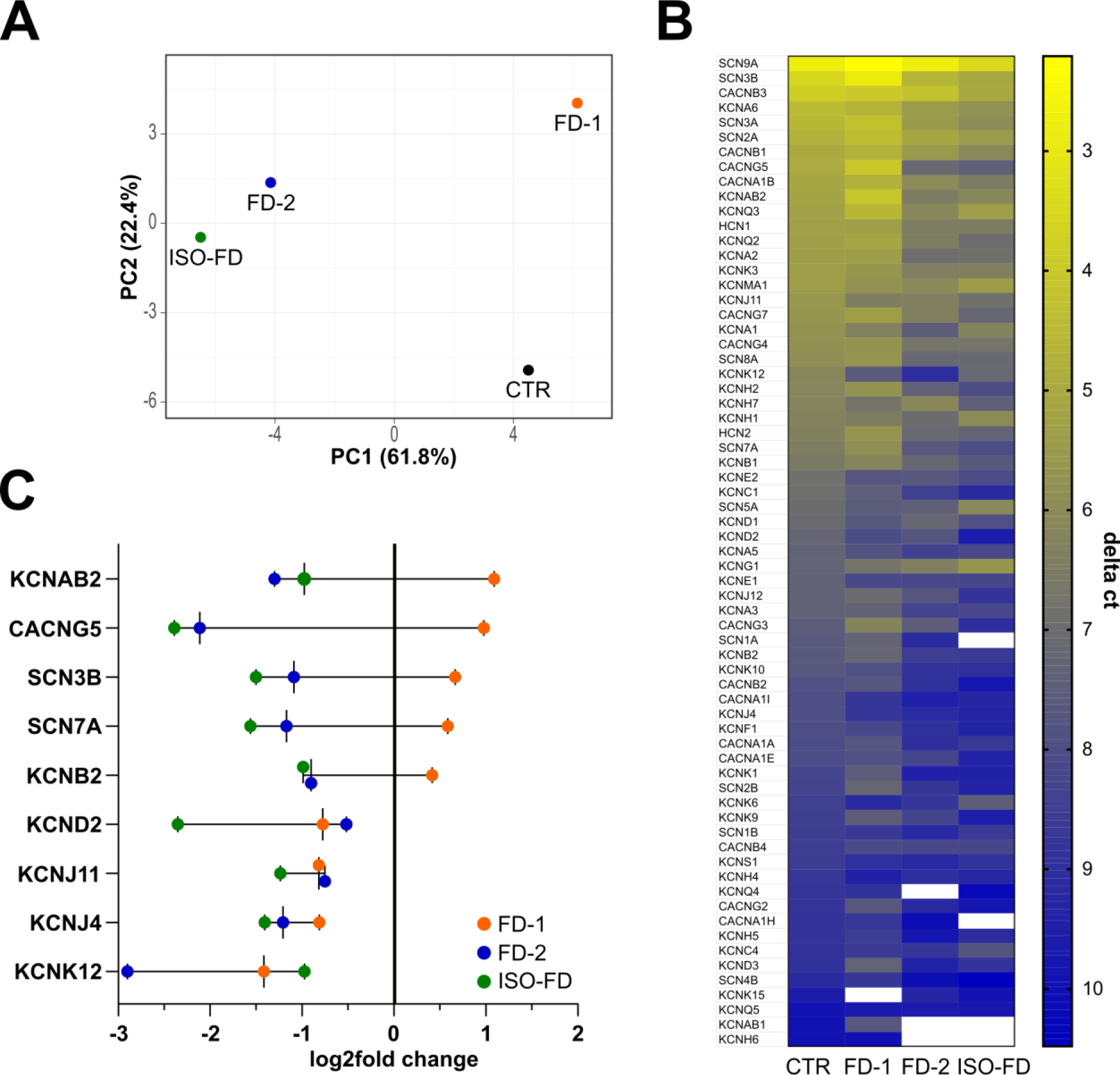
Ion channel gene expression of sensory neurons. (**A**) Principle component analysis of voltage-gated ion channel array expression. **(B)** Heatmap of expressed (C_t CTR_ < 33) genes from highest (yellow) to lowest (dark blue) expression indicated as ΔC_t_. **(C)** Exemplary gene transcripts illustrating the inverse regulation between FD-1 versus FD-2 and ISO-FD and a subgroup of unanimously downregulated genes compared to Ctrl baseline. **Abbreviations:** Ctrl = control, FD-1, FD-2 = patients with Fabry disease, ISO-FD = isogenic Fabry line, PCA = principal component analysis.

### Sensory neurons of FD patients exhibit temperature-dependent hypoexcitability

For functional analysis of sensory neurons, we next performed patch-clamp recordings at RT and 39°C as surrogate for clinical fever. Baseline electrophysiological characteristics of the investigated sensory neurons gave similar profiles in all cell lines (Fig. 5A - C, Supplementary Fig. 5A - D). Exposure of neurons to 39°C resulted in increased firing frequencies of action potentials without inter-individual differences (Fig. 5A; Supplementary Fig. 5E). Accordingly, shortening of action potential duration occurred at 39° as indicated by decreased half-widths (p < 0.0001; Fig. 5B) and increased rising slopes (p < 0.0001; Fig. 5C).

**Figure 5:**
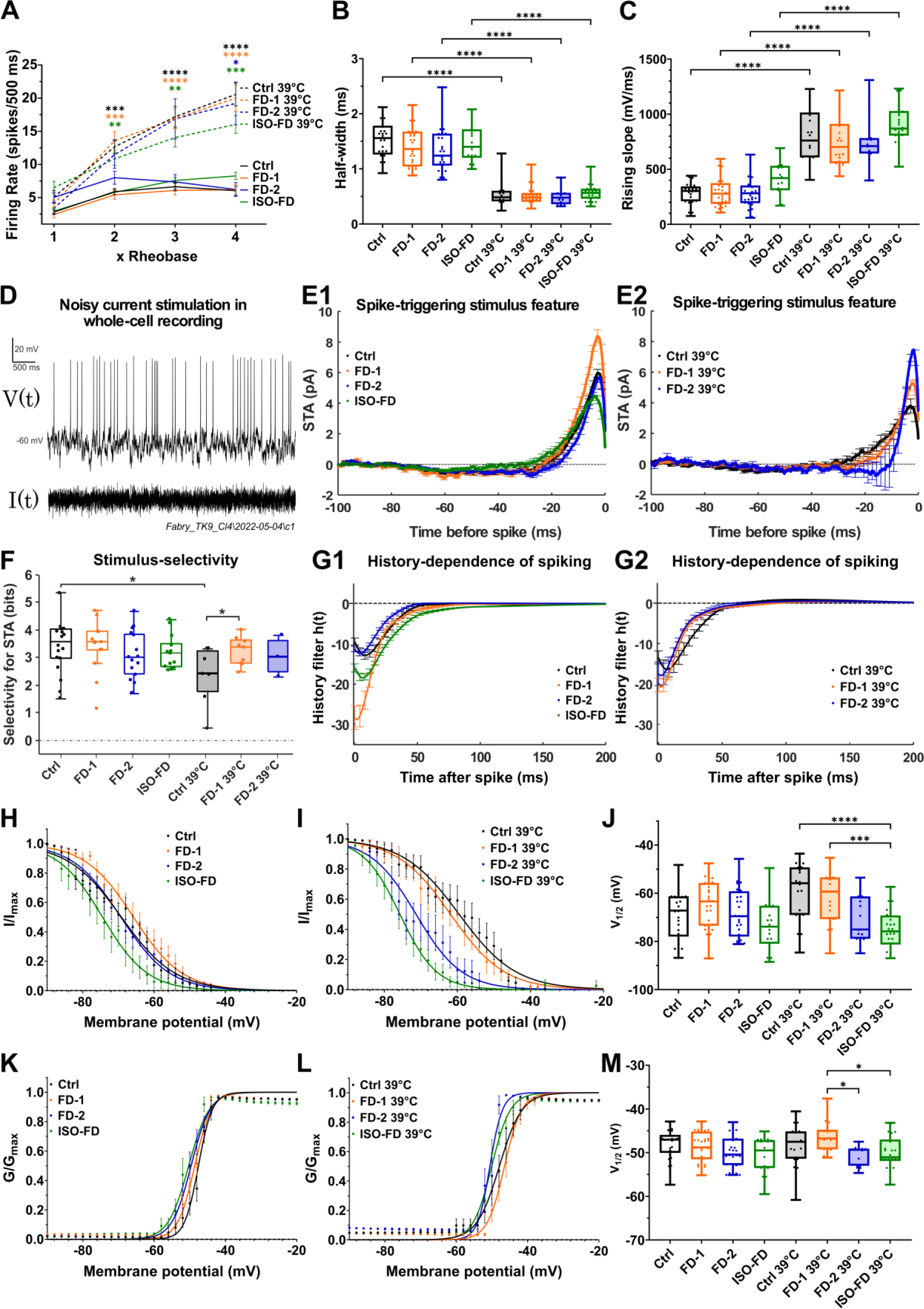
Electrophysiological characterisation of sensory neurons. (**A**) No differences were found in action potential firing rates upon injection of 1x, 2x, 3x, and 4x rheobase current between cell lines. Increased firing rates were observed at 39°C compared to RT at 2x and 3x rheobase for Ctrl (n = 2 clones; RT: clone 1 = 7 cells, clone 2 = 21 cells; 39°C: clone 1 = 16 cells, clone 2 = 4 cells), FD-1 (n = 2 clones; RT: clone 1 = 11 cells, clone 2 = 16 cells; 39°C: clone 1 = 18 cells, clone 2 = 5 cells), and ISO-FD (n = 1 clone; RT: 19 cells, 39°C: 26 cells) neurons. At 4x rheobase, all cell lines (FD-2: n = 2 clones; RT: clone 1 = 5 cells, clone 2 = 18 cells; 39°C: clone 1 = 15 cells) showed increased firing rates at 39°C compared to RT. Data are represented as mean ± SEM. Two-way ANOVA followed by Sidak’s multiple comparison correction. (**B**) Half-width of action potentials was comparable between Ctrl, FD-1, FD-2, and ISO-FD neurons at RT and 39°C. Reduced half-widths at 39°C for all cell lines compared to RT. (**C**) Rising slope of action potentials was not different between Ctrl, FD-1, FD-2, and ISO-FD neurons at RT and 39°C. Increased rising slopes at 39°C for all cell lines compared to RT. (**D**) Neurons were stimulated with Gaussian white current in patch-clamp whole-cell recordings: voltage response (upper) and noisy stimulus (lower). Representative recording shown. Group sizes for RT and (39°C) were n = 18 (7), 14 (9), 15 (3), 13 (4) for Ctrl, FD-1, FD-2, and ISO-FD, respectively. (**E**) STA illustrates mean current eliciting action potentials calculated via spike-triggered reverse correlation. Positive values indicate depolarizing current; t = 0 indicates the time of spiking. At RT, FD-1 neurons required a larger depolarizing STA compared to the other cell lines (E1). At 39°C, both FD cell lines encoded larger depolarizations compared to Ctrl (E2). Data are represented as mean ± SEM. (**F**) Stimulus selectivity for the STAs shown in **E1,2** measured in bits was comparable between the cell lines at RT, but increased in FD-1 at 39°C. Large values indicate high selectivity for the STA; i.e. the spike-triggering subspace defined by the STA is very different from the overall Gaussian stimulus distribution. Rank-sum test. (**G**) History-dependence of neuron populations, *h(t)*, calculated via Generalized Linear Model framework (see Methods) showed a higher refractoriness of FD-1 compared to the other cell lines at RT (G1), which largely disappeared at 39°C (G2). Data are represented as mean ± SEM. (**H**) Steady-state inactivation curves of voltage-gated sodium channels at RT. ISO-FD neurons showed negative shift of inactivation compared to Ctrl (V_1/2_: Ctrl: −69.18 mV; FD-1: −65.79 mV; FD-2: −69.32 mV; ISO-FD: −74.46 mV). Data are represented as mean ± SEM. (**I**) Steady-state inactivation curves of voltage-gated sodium channels at 39°C. FD-2 and ISO-FD neurons displayed negative shift of inactivation compared to Ctrl (V_1/2_: Ctrl: −59.51 mV; FD-1: −61.86 mV; FD-2: −71.32 mV; ISO-FD: −75.73 mV). Data are represented as mean ± SEM. (**J**) Comparison of V_1/2_ steady-state inactivation. V_1/2_ was comparable between Ctrl, FD-1, FD-2 and ISO-FD neurons at RT. However, at 39°C, V_1/2_ was decreased in ISO-FD and FD-2 showed a trend towards decreased V_1/2_ compared to Ctrl and FD-1. (**K**) Steady-state activation curves of voltage-gated sodium channels at RT were comparable between the cell lines at RT (V_1/2_: Ctrl: −47.57 mV; FD-1: - 48.62 mV; FD-2: −49.64 mV; ISO-FD: −50.3 mV). Data are represented as mean ± SEM. (**L**) Steady-state activation curves of voltage-gated sodium channels at 39°C displayed a negative shift of FD-2 and ISO-FD compared to Ctrl and FD-1 (V_1/2_: Ctrl: −47.9 mV; FD-1: −46.47 mV; FD-2: −50.46 mV; ISO-FD: −50.4 mV). Data are represented as mean ± SEM. (**M**) Comparison of V_1/2_ steady-state activation. V_1/2_ was comparable between Ctrl, FD-1, FD-2, and ISO-FD neurons at RT. At 39°C, V_1/2_ showed a trend towards decreased values for FD-2 and ISO-FD compared to Ctrl. For (B-C, H-M): For Ctrl (n = 2 clones; RT: clone 1 = 8 cells, clone 2 = 21 cells; 39°C: clone 1 = 16 cells, clone 2 = 4 cells), FD-1 (n = 2 clones; RT: clone 1 = 15 cells, clone 2 = 16 cells; 39°C: clone 1 = 18 cells, clone 2 = 5 cells), FD-2 (n = 2 clones; RT: clone 1 = 11 cells, clone 2 = 18 cells; 39°C: clone 1 = 15 cells), and ISO-FD (n = 1 clone; RT: 19 cells, 39°C: 26 cells) pooled data obtained from ≥ 3 individual differentiations were used. For (D-G): Group sizes for RT and (39°C) were n=18 (7),14 (9),15 (3),13 (4) for Ctrl, FD-1, FD-2, and ISO-FD, respectively. Data were pooled from 2 clones per cell line (exc. ISO-FD). For (B-C, F, J, M): Data are represented as box-and-whisker plots with dots as individual values. The box width indicates the first and third quartiles, the line indicates the median, and the whiskers of the box plot indicate the smallest and largest values. For (B-C, J, M): One-way ANOVA followed by Sidak’s multiple comparison correction. **Abbreviations:** STA = spike-triggered average stimuli, Ctrl = control, RT = room temperature, FD-1, FD-2 = patients with Fabry disease, ISO-FD = isogenic Fabry line. p < 0.05 = *; p < 0.01 = **; p < 0.001 = ***; p < 0.0001 = ****.

We next asked whether the cell lines systematically differed in ability to encode time-varying stimuli using Linear-Nonlinear and Generalized Linear point process (GLM) models. Action potentials were evoked with a noisy current stimulus (Fig. 5D, Supplementary Methods) and spike-triggered average (STA) currents (Fig. 5E), selectivity for the STA (Fig. 5F), and spike-history effects quantified via GLM analysis were compared (Fig. 5G). All sampled neurons were effectively driven by noisy current patterns and typically integrated current within a 50-ms window. Furthermore, both stimulus and history encoding were markedly temperature dependent.

At RT, the FD-1 population required a larger depolarizing STA (Fig. 5E1) relative to the other populations. However, at 39°C, both FD cell lines encoded larger depolarizations relative to Ctrl (Fig. 5E2). Stimulus selectivity, i.e. how precisely spike-evoking stimuli matched the STA, was similar across groups measured at RT, but more variable at 39°C (Fig. 5F). Most strikingly, at 39°C, the FD-1 group was more selective for the STA relative to Ctrl (p < 0.05). Moreover, GLM analysis of statistical interactions between spikes revealed that the FD-1 population was ∼2-3x more refractory than the other populations at RT (Fig. 5G1), while these differences largely disappeared at 39°C (Fig. 5G2).

These data suggest that alterations in both stimulus- and history-encoding properties in FD-1 neurons support a functional decrease in excitability and a sensitivity to temperature, potentially contributing to the clinical hyposensitivity to thermal stimuli determined by QST. In contrast, the FD-2 population showed encoding properties more similar to that of Ctrl.

We then asked whether voltage-gated sodium channels are involved in the reduced excitability of FD-1 neurons. While sodium current densities were largely unaffected (Supplementary Fig. 5F-I), investigation of steady-state inactivation curves showed a moderately hyperpolarized shift of inactivation of ISO-FD at RT (Fig. 5H). At 39°C, the negative shift of fast inactivation was even more pronounced in ISO-FD neurons and, to a lesser extent, also present in FD-2 neurons (Fig. 5I). This finding is in accordance with decreased sodium current availability due to steady state inactivation. Similarly, V_1/2_ steady-state inactivation was decreased in ISO-FD at 39°C (p < 0.0001; Fig. 5J), whereas V_1/2_ in FD-2 was not different from Ctrl. Steady-state activation curves were comparable between all cell lines at RT (Fig. 5K). At 39°C, steady-state activation of FD-2 and ISO-FD was slightly shifted towards hyperpolarized potentials compared to Ctrl and FD-1 (Fig. 5L). However, V_1/2_ steady-state activation for FD-2 and ISO-FD was similar compared to Ctrl and decreased compared to FD-1 (p < 0.05; Fig. 5M). Steady-state activation curves of voltage-gated potassium channels did not show any differences between the cell lines (Supplementary Fig. 5J-K)

### Heat increases Gb3-dependent neuronal Ca^2+^ levels and thinning of neurite calibre

We next used confocal Ca^2+^ imaging under physiological (37°C) and clinical fever (39°C; Fig. 6A) conditions and found elevated neuronal Ca^2+^ levels in FD-1 and FD-2 neurons compared to Ctrl neurons already at 37°C (p < 0.001; Fig. 6B). At 39°C, Ca^2+^ concentrations further increased dramatically in neurons of both FD cell lines compared to 37°C and to Ctrl neurons (p < 0.001 each; Fig. 6B). Interestingly, neurite calibres of FD-1 and FD-2 showed substantial thinning compared to those of Ctrl neurons at baseline (p < 0.001; Fig. 6C) and only neurites of FD-1, who reported typical heat-triggered Fabry pain, displayed further thinning upon heat stimulation (p<0.001; Fig. 6C). Neuronal identity was assured by final KCl application leading to excessive Ca^2+^ activity (Supplementary Video S1).

**Figure 6:**
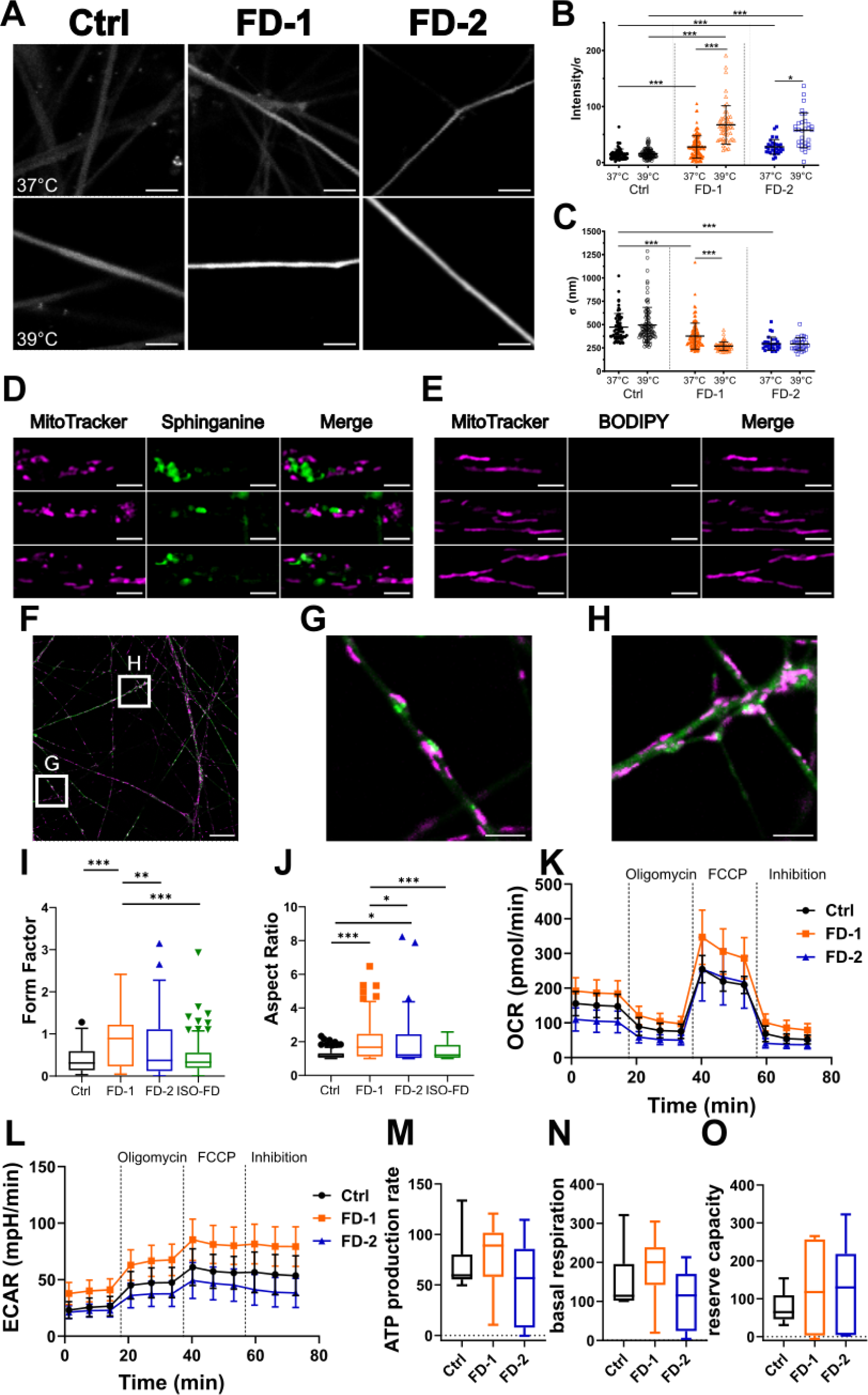
Sensory neuron heat stimulation and mitochondrial characteristics. **(A)** Exemplified micrographs from firing neurons at 37°C (upper row) and 39°C (lower row). Scale bars: 5 µm. **(B)** FD-1 and FD-2 neurons show higher activity at 37°C (FD-1: n=1 clone, 179 neurites; FD-2: n=1 clone, 30 neurites) and increased calcium signalling after incubation at 39°C (FD-1: n=1 clone; 56 neurites; FD-2: n=1 clone, 31 neurites), whereas the Ctrl did not show an increase upon heat (37°C: n=1 clone, 63 neurites; 39°C: n=1clone, 103 neurites). Data are represented as mean ± SD. Kruskal-Wallis test with Bonferroni correction for multiple testing. **(C)** FD-1 and FD-2 neurons show a generalized decrease in neurite diameter and FD-1 shows further thinning upon heat stimulation. N=equal to (B). Data are represented as mean ± SD. Kruskal-Wallis test with Bonferroni correction for multiple testing. **(D)** Metabolic labelling with sphinganine in Ctrl neurons reveals mitochondrial fragmentation in the vicinity of accumulations. Scale bars: 2 µm. **(E)** In contrast, incubation of Ctrl neurons with BODIPY only, shows normal mitochondrial morphology. Scale bars: 2 µm. **(F)** Still image of mitochondria tracking of FD-1 neurons. Scale bar: 25 µm. **(G)** Still image of mitochondria/sphinganine interaction. Scale bar: 5 µm. **(H)** Still image of potential mitochondrial block by sphinganine. Scale bar: 5 µm. (G) + (H) Cropped from (F). **(I)** FD-1 neurons show mitochondria with an increased form factor compared to all other cell lines. **(J)** FD-1 and FD-2 neuronal mitochondria show an increased aspect ratio compared to Ctrl. Aspect ratio of FD1 neuronal mitochondria is increased compared to FD-2. For (I) + (J): pooled data from n = 2 clones/line, obtained from 3 individual differentiations each was used (20 photomicrographs per differentiation = 60 per clone were analysed). Data are represented as Tukey boxplot. Kruskal-Wallis test with Dunn’s multiple comparison was applied. **(K) + (L)** Comparison of OCR and ECAR of Ctrl and FD neurons using Seahorse assay showed no difference in cellular metabolism. Data are represented as mean ± SEM. Two-way ANOVA with multiple comparison. Seahorse assay also showed no difference in ATP production **(M)**, basal respiration **(N)** and reserve capacity **(O)** between Ctrl and Fabry patients. Data are represented as Tukey boxplot. Kruskal-Wallis test with Dunn’s multiple comparison was applied. Seahorse assay was performed from 3 individual differentiations of n = 2 clones/line. **Abbreviations:** Ctrl = control, FD-1, FD-2 = patients with Fabry disease, ISO-FD = isogenic Fabry line, OCR= oxygen consumption rate, ECAR= extracellular acidification rate. p < 0.05 = *; **p < 0.01 = **; p < 0.001 = ***.

### Fabry sensory neurites exhibit mitochondrial traffic jam and altered morphology

While Gb3 can be visualized in fixed cells applying STxB, direct investigation during live-cell imaging was not possible. Hence, we used a metabolic, click-chemistry-based labelling approach with the Gb3 precursor ω-N_3_-sphinganine (sphinganine).^37^ Qualitatively, we observed mitochondrial fragmentation and shrinkage in FD sensory neurites mainly in close vicinity to sphinganine accumulations (Fig. 6D) which were absent in dye-control samples (Fig. 6E). In FD-1 neurons, we further observed jam of mitochondrial trafficking at sphingolipid accumulations (Fig. 6F - 6H; Supplementary Videos S2 – S4).

Quantitatively, mitochondria in FD-1 neurons were more branched compared to FD-2, ISO-FD, and Ctrl (p < 0.01; Fig. 6I) and the aspect ratio was higher in FD cell lines compared to Ctrl (p < 0.05; Fig. 6J). Investigation of mitochondrial functionality using Seahorse assays did not show any differences between FD and Ctrl lines regarding oxygen consumption and extracellular acidification rate (Fig. 6K – 6L). Basal respiration, ATPase-dependent respiration, and reserve capacity were also not altered in FD neurons (Fig. 6M – 6O).

## DISCUSSION

We pioneer *in vitro* evidence linking GLA impairment with small fibre neuropathy in Fabry disease as one of its major clinical hallmarks. We show that patient-derived sensory neurons are activated by heat, electrically hypoexcitable, and show hints for mitochondrial traffic jam within neurites which may contribute to triggerable neuropathic pain, thermal hyposensitivity, and denervation in Fabry disease.

*In vitro* models have been generated for genetic pain syndromes using iPSC-derived sensory neurons^15–17,44^ and have helped to unravel disease pathophysiology^15,44,45^ and identify novel targets for treatment.^16,17^ In Fabry research, generation of sensory neurons retaining cellular disease phenotype using patient biomaterial was not successful so far,^46,47^ while differentiation of Fabry iPSC to cardiomyocytes,^11,48^ podocytes,^49^ or vascular endothelial cells^50^ was achieved. We modified a published differentiation strategy^32^ and generated iPSC-derived sensory neurons showing pathognomonic Gb3 accumulations.

Pain in Fabry disease is one of the very early symptoms starting in childhood^51^ and is mostly episodic and triggerable.^4,52^ Using our *in vitro* disease model, we show that heat, a typical trigger of Fabry pain, leads to increased Ca^2+^ levels in Fabry neurons, pointing to higher neuronal activity (Fig. 6B). A potential link between Fabry pain and Ca^2+^ levels was already assumed: studies reported alterations of Ca^2+^-activated ion channel expression in Fabry disease, such as K_Ca_1.1 in patient fibroblasts,^53^ K_Ca_3.1 in Gb3-treated human umbilical vein endothelial cells, and aortic endothelial cells from a Fabry mouse model.^54^ Further, elevated intracellular Ca^2+^ concentrations were associated with increased lyso-Gb3 levels, the deacylated derivative of Gb3^55^ in murine DRG neurons leading to pain-related behavior.^10^

We extended knowledge on sensory neuron ion channel RNA expression profiles and found that distinct ion channels were inversely expressed between FD-1 versus FD-2 and ISO-FD neurons. Among these, KCNAB2 was shown to promote TRPV1 activity,^56^ while SCN3B and SCN7A were positively correlated with neuropathic or bone pain in animal models.^57,58^ In contrast, KCNB2 can lead to hyperexcitability when downregulated.^59^ Overall, FD-1 sensory neurons may be tuned towards a more pain promoting expression profile, matching patients’ phenotype, although potential methodological limitations such as variations in the amount of sensory neuron sub-populations between differentiations and clones preclude more detailed conclusions. Further, several voltage gated potassium channel family members were collectively downregulated in all FD lines and may hint to a partially conserved disease-specific channel expression pattern. Interestingly, Kir6.2 deficiency, encoded by KCNJ11, resulted in small fibre dysfunction and axonal degradation in mice.^60^

We found thinner neurites of Fabry sensory neurons compared to those of control neurons. Fibre thinning was progressive upon heat simulation exclusively in neurons of FD-1, who also reported pain. Although literature on neurite diameter is sparse, one potential cause for temperature-dependent neuronal shrinkage is hypoxia as shown in ischemic brains of squirrel monkeys.^61^ We hypothesize that heat leads to increase in neuronal reactive oxygen species in Fabry disease^50,62^ depleting intracellular oxygen with consecutive cellular shrinkage. Little is known about pain-associated reduction of nerve fibre diameters in the peripheral nervous system, however, there are examples of reduced intraepidermal nerve fibre calibres e.g. in fibromyalgia syndrome.^63^ Also, corneal nerve fibre diameter was reduced in patients with small fibre pathology^64^ correlating with disease severity.^65^ While the exact mechanism remains to be elucidated, reduction of neuronal membrane surface leads to reduced cell capacitance and higher dendritic length constant. This, in turn, facilitates signal propagation and AP firing,^66^ which may be a contributor to the characteristic heat-triggered Fabry pain phenotype.

Confocal Ca^2+^ imaging enabled the investigation of bulk Ca^2+^ levels in a single focal plane with subcellular resolution, but at the cost of temporal resolution. This technical limitation might mask subtle differences in AP frequency, or other single AP related parameters. Still, we show that heat exclusively increases Ca^2+^ levels of Fabry-derived sensory neurons, but spares Ctrl neurons, mimicking the triggerable aspect of clinical Fabry pain by fever. The lack of an increased Ca^2+^ signal in Ctrl neurons upon heat stress strongly suggests that Gb3 is crucially involved in this process.

Age-dependent thermal hyposensitivity is a major symptom particularly in men with Fabry disease. Systematic analysis of warm and cold detection thresholds revealed progressive elevation of perception thresholds even under continued enzyme replacement therapy (ERT).^67^ Studies investigating the *GLA* KO mouse model of Fabry disease showed analogous findings.^68,69^ Electrophysiological recordings gave evidence for reduced sodium current densities in the *GLA* KO mouse model as potentially underlying mechanism, which was Gb3- and age-dependent.^6^ Using extensive electrophysiological analysis including GLM analysis on iPSC-derived sensory neurons of Fabry patients, we now show a negative shift of fast steady state inactivation of voltage-gated sodium channels and FD neurons also needed larger spike-triggered average current in accordance with higher inactivation (Fig. 5). It is known that a negative fast inactivation shift leads to pain alleviation,^70^ a common mechanism utilized in Na_v_ channel blocker-based analgesics,^71^ which makes our finding highly interesting for potential druggable targets in analgesic treatment of Fabry pain. In an *in vitro GLA* KO model, we showed that Gb3 directly reduces Na_v_ 1.7 current densities, which can be rescued by AGAL treatment.^6^ We speculate that elevated inactivation of sodium channels may underlie reduced warm and cold perception typically found in Fabry patients. To decipher the subcellular mechanism linking Gb3 deposits with the observed functional alterations in voltage-dependent sodium channels, further studies are needed. In our previous study,^6^ we showed that Na_v_1.7 electric properties were altered in *GLA* KO mice, although its protein expression was unaffected. Therefore, future investigations using our human *in vitro* system should not only focus on expression profiles, but also on membrane localization and cytosolic transport of pain-relevant ion channels. While the cellular function of the membrane lipid Gb3 is still unknown, its mere increase might already account for membrane disturbance as was shown in Fabry fibroblasts.^72^ Hence, an impact of Gb3 on channel clustering^73^ or membrane anchoring could also induce the detected abnormalities.

In contrast to our *in vitro* findings, continued ERT does not improve thermal detection thresholds in Fabry patients,^67^ which may be explained by a better response of rejuvenated cells during iPSC generation compared to adult cells *in vivo* with chronic Gb3 overload.^74,75^ Besides iPSC-specific effects, it is unclear if and to which extent ERT crosses the blood-nerve-barrier *in vivo.*^76^ In our *in vitro* model, nociceptors lack the components typically found in the blood-nerve-barrier, such as endoneurial endothelial cells, which limit the transport of proteins into the DRG.^77^ Instead, AGAL can directly enter the neuron, likely via the mannose-6-phosphate receptor.^78^ Analysis of tissue from a deceased Fabry patient revealed that Gb3 accumulations were abundant in all organ systems including the DRG despite ERT.^79^ This further hints to an insufficient permeability of the blood-nerve-barrier for ERT, potentially explaining the differences between the *in vivo* and *in vitro* situation.

Another characteristic of Fabry disease is peripheral denervation as reflected by reduced IENFD in skin punch biopsies.^67,80^ Using metabolic labelling, we observed a mitochondrial traffic jam at sphinganine deposits in neurites, which may contribute to dying-back peripheral denervation. We further found mitochondrial dysmorphism in Fabry sensory neurons. There is growing evidence for impaired mitochondrial function in lysosomal storage disorders including Fabry disease.^81^ Mitochondrial dysfunction already was linked to neurodegenerative diseases such as Alzheimer’s disease,^82^ Parkinson’s disease^83^ or amyotrophic lateral sclerosis.^84^ Quantitative metabolic function was not different between FD sensory neurons, as was recently also shown for podocytes.^62^ Still, our finding of altered mitochondrial morphology is in line with a recent study reporting glucosylceramide accumulations in murine dopaminergic neurons that trigger impaired interaction of mitochondria and lysosomes, and mitochondrion depletion in neurites.^85^

We present a patient-specific neuronal and functional human *in vitro* disease model for Fabry disease providing several crucial findings on Fabry pathophysiology: 1) Gb3 accumulates ubiquitously in sensory Fabry neuronal somas and neurites. 2) Lysosomal integrity is impaired displaying a high Gb3 load potentially contributing to fibre degeneration. 3) Pain-related voltage-gated sodium channels from Fabry patients show a differential inactivation kinetic. 4) Neuronal Gb3 accumulations lead to heat-induced Ca^2+^ increase and a decrease in neuronal diameter as potential basis of Fabry pain. 5) Sphingolipid accumulations impair mitochondrial dynamics and morphology, and may underlie nerve fibre degeneration. Our *in vitro* model opens the avenue to study patient-specific disease mechanisms in a multi-dimensional approach, thus, paving the way towards the development of targeted treatments not only acting on the deficient enzyme itself, but also preventing cellular defects as a result of increased neuronal Gb3 load in patients with Fabry disease.

## Supporting information

Supplementary Meterial

## Acknowledgements

We thank Daniela Urlaub, Danilo Prtvar, Kathleen Stahl (Department of Neurology, University of Würzburg, Germany), and Birgit Halliger-Keller (Institute for Human Genetics, University of Würzburg, Germany) for expert technical assistance. We further thank our undergraduate students Frederik Bär, Viktoria Diesendorf, Lisa Henkel, Hannah Kiefer, Philipp Neundorf, and Maxine Sokolowski (Department of Neurology, University of Würzburg, Germany) for help during cell cultivation.

