## Supplementary Meterial for "Small fibre neuropathy in Fabry disease: a human-derived neuronal *in vitro* disease model"

### Supplementary Material

#### Methods

##### Immunoreactions

For ICC, cells were fixed with 4% paraformaldehyde (PFA; Electron Microscopy Sciences, Hatfield, USA) in $PBS^{++}$ (Merck, Darmstadt, Germany), or 4% PFA in Live Cell Imaging Solution (Thermo Fisher Scientific, Waltham, USA) for neurons, respectively, for 15 min at room temperature (RT). Cells were washed three times with $PBS^{++}$ and blocked for 30 min at RT with 10% FCS + 0.1% saponin (Sigma-Aldrich, St. Louis, MO, USA) for intracellular targets, without saponin for surface markers. Primary antibodies were diluted in blocking solution and incubated for 16 h at 4°C and washed afterwards. Secondary antibodies were diluted in $PBS^{++}$ and incubated for 2 h at RT (30 min for TOM20), nuclei were counterstained with 700 nM 4',6-diamidino-2-phenylindole (DAPI; Sigma-Aldrich, St. Louis, MO, USA), washed three times with $PBS^{++}$ and mounted using ProLong Glass antifade mountant (Thermo Fisher Scientific, Waltham, MA, USA) for super-resolution microscopy and Aqua-Poly/Mount (Polysciences, Warrington, PA, USA) for ICC. Five µg/ml StxB (Sigma Aldrich, St. Louis, MO, USA) was conjugated with Alexa Fluor 555 (all: Thermo Fisher Scientific, Waltham, MA, USA) or Atto 643 (for ExM; ATTO-TEC, Siegen, Germany) using Zeba spin desalting columns (Thermo Fisher Scientific, Waltham, MA, USA). Sixteen ng/ml StxB::555 was added during the secondary antibody incubation, where applicable.

For immunohistochemistry, cryosections fixed in PFA and embedded in Tissue-Tek O.C.T. (Sakura, Osaka, Japan) were thawed, blocked with 10% bovine serum albumin (BSA) in PBS + 0.1% saponin and immunoreacted with the primary antibody (PGP9.5) or STxB::555

diluted in 1% BSA/PBS + 0.1% saponin overnight at RT. The next day, sections were washed and incubated with the corresponding dye-conjugated secondary antibody, where applicable. Sections were mounted using VECTASHIELD with DAPI (Vector Laboratories, Burlingame, CA, USA) and sealed with Covergrip Coverslip sealant (Biotium, Fremont, CA, USA).

#### Expansion microscopy

For expansion microscopy (ExM),[1,2] cells were labelled and imaged prior to expansion with an inverse microscope (DMi8, Leica, Wetzlar, Germany). Cells were incubated with 0.25% glutaraldehyde (SERVA, Heidelberg, Germany) in PBS for 10 min at RT. Gelation was performed with monomer solution (8.625% sodium acrylate, 2.5% acrylamide, 0.15% N,N‘-methylenbisacrylamide, 2 M NaCl in PBS, mixed with 0.2% ammonium persulfate, and 0.2% tetramethylethylenediamine; all Sigma-Aldrich, St. Louis, MO, USA) for 30 min at RT. Gels were incubated for 30 min at 37°C in digestion buffer containing 50 mM Tris, pH 8.0 (SERVA, Heidelberg, Germany), 25 mM EDTA (Thermo Fisher Scientific, Waltham, MA, USA), 0.5% Triton X-100 (Sigma-Aldrich, St. Louis, MO, USA ), 0.8 M NaCl (Th. Geyer, Renningen, Germany), and 4 U/ml proteinase K (NEB, Ipswich, MA, USA). Gels were washed once for 10 min at RT with PBS, followed by 700 nM DAPI (for iPSC: 10 µM Nuclear Orange [AAT Bioquest, Sunnyvale, CA, USA] was added) in PBS for 20 min, and again only PBS for 10 min. Expansion was done in excess dd$H_2O$ for 1 h at RT. Gels were transferred into poly-D lysine (Sigma Aldrich, St. Louis, MO, USA) coated chambers (Thermo Fisher Scientific, Waltham, MA, USA). Expansion factor was determined using nuclear and lysosomal associated membrane protein 1 (LAMP1) signal of same regions via a published automated python script[3] or manual alignment in Inkscape (version 0.92.3, 11.03.2018). Imaging was performed using a Zeiss

LSM700 for iPSC, or a Zeiss Elyra 7 for neurons (both: Zeiss, Oberkochen, Germany).

#### Patch-clamp analysis

##### Whole-cell

Whole-cell patch-clamp analysis was performed on iPSC-derived neurons between five and eight weeks after differentiation. Currents were recorded at RT or at 39°C using a warmed platform (WP-10, Warner Instruments, Holliston, MA, USA) connected to a single channel heater controller (TC-324C, Warner Instruments, Holliston, MA, USA). Bath solution consisted of 180 mM NaCl, 5.4 mM KCl, 1.8 mM $CaCl_2$, 1 mM $MgCl_2$, 10 mM glucose, and 5 mM HEPES.[4,5] Patch pipettes were pulled from borosilicate glass capillaries (Kimble Chase Life Science and Research Products, Meiningen, Germany) and were heat-polished to reach a pipette resistance of 3 to 6 MΩ (whole-cell). The pipette recording solution contained 170 mM KCl, 2 mM $MgCl_2$, 1 mM EGTA, 1 mM ATP, and 5 mM HEPES. In both, bath, and pipette solution, osmolarity was adjusted to 280-300 mOsm. Currents were recorded with an EPC10 patch-clamp amplifier (HEKA, Ludwigshafen, Germany) with a sampling rate of 20 kHz. Stimulation and data acquisition were controlled by the Patchmaster software package version v2x90.5 (HEKA, Lambrecht, Germany) on a windows computer, and data analysis was performed off-line with GraphPad PRISM version 9.5.0 (GraphPad Software, Inc., La Jolla, CA, USA) and OriginPro 2021b version 9.8.5.212 (OriginLab Corporation, Northampton, MA, USA). Neurons with the ability to elicit more than one action potential (AP) were considered nociceptors.

##### Current clamp

APs were induced by square current injections (from 0 to 290 pA in 10 pA steps). The first AP evoked was used to calculate the AP parameters. Analysis of APs was performed with

Stimfit version 0.15.8.[6] The threshold potential was measured at the point of inflection during the AP depolarizing phase characterized by a slope of 10 mV/ms. To evaluate AP amplitude, the difference between membrane potential and AP peak was used. Rise and decay slopes were measured from 20% to 80% of peak amplitude. AP duration was calculated by measuring the distance between depolarization and repolarization at the point of half maximum AP amplitude. Rheobase was defined as the minimum current required to elicit an AP. Repetitive firing of action potentials was analysed by counting the number of AP upon injection of 1x, 2x, 3x and 4x rheobase current in 500 ms steps.

**Voltage clamp**

Voltage-gated sodium ($Na_V$) current densities were obtained by normalizing $Na_v$ current amplitudes to the cell capacitance. Outward voltage-gated potassium ($K_v$) currents were recorded with the same voltage-step protocol and $K_v$ current amplitudes were normalized to the cell capacitance to receive $K_v$ current densities. To acquire current-voltage curves, neurons were held at membrane potential for 20 ms followed by a 50 ms prepulse at -100 mV. Voltage pulses from -90 mV to +8 mV were applied in 2 mV steps for 200 ms. After depolarization steps, neurons were held at maximum $Na_V$ activation potential for 200 ms to measure $Na_V$ channel inactivation kinetics. Normalized conductance-voltage curves were fitted with a Boltzmann equation:

$$Y = 1/(1 + exp\ [(V\ - V_{50})/k]). \qquad (1)$$

Y is G/Gmax, V is the membrane voltage, $V_{50}$ is the voltage at half-maximal channel activation, and k is the slope factor.

#### Linear-Nonlinear and Generalized Linear models

To characterize the transformation from current to action potentials, Linear-Nonlinear (LN) cascade and Generalized Linear point process models (GLM) were calculated for each cell using spike times evoked by current-clamp stimulation with broad spectrum (near-white, exponentially filtered with time constant $\tau$=1 ms) Gaussian noise, $I(t)$, as has been described for single neurons elsewhere;[7,8] see here[9] for general review of the methods.

For a simple LN characterization of excitability, we calculated $f$, the spike-triggered average (STA) current via spike-triggered reverse correlation, and projected the raw stimulus $I(t)$ into the dimension defined by $f$ via convolution:

$$s(t) = f * I(t). \qquad (2)$$

To quantify sensitivity to this feature in bits of information, the distribution of spike-triggering stimuli, $P(s|sp)$, was compared to the overall filtered stimulus distribution, $P(s)$, using the Jensen-Shannon divergence, $DJS$, as has been described previously.[7,8] Large $DJS$ values indicate strong selectivity for $f$ and thus very specific encoding of the stimulus feature, whereas $DJS$ values close to zero indicate little selectivity. For each neuron, $DJS$ was calculated as the average of 100 bootstrap resamples from $P(s|sp)$.

The simple formulation of the LN model described above does not account for interspike interactions. To capture both stimulus-driven spiking and non-Poisson spiking effects, GLM models were fit to the same data set. In brief, GLM methods expand upon the LN framework to simultaneously fit a stimulus filter $f(t)$ and an autoregressive spiking-history-dependent filter, $h(t)$. $h(t)$ transiently modulates the probability of spiking for a period after a spike predicted by large values of $f(t)$, thereby taking past spiking history into account. Both filters are fit by negative log-likelihood minimization. To lower the dimensionality of parameter space to be fitted, a raised cosine basis vectors approach was used.[10,11] We used five basis vectors for each filter, and filter length t = 100 ms and t = 275 ms for $f(t)$ and $h(t)$, respectively. In the data set analyzed here, the GLM stimulus filters were largely similar to the STA calculated directed from the data, so only $f(t)$ is presented in Figure 5G.

All analysis and stimulus generation was done in Matlab 2019b or 2022b (The MathWorks Inc., Natick, MA, USA). Only neurons with stable resting membrane potentials were included. Spike times were identified by examination of dV/dt vs. V and finding inflection points in the first derivative dV/dt, using a user-defined threshold for each cell. 547-3037 (mean 1436 ± 495) spikes were used for LN and GLM analysis; stable GLM filters were obtained for data sets including more than 300 spike times.

#### X-chromosome inactivation analysis and *GLA* transcription

For *GLA* expression analysis, a PCR of cDNA from FD-3 iPSC was performed using primers flanking the *GLA* mutation site. PCR products were purified and Sanger sequenced (Eurofins Genomics, Ebersberg, Germany). For X-chromosome inactivation (XCI) analysis, DNA from FD-3 iPSC was isolated using a commercial kit (DNeasy Blood & Tissue; Qiagen, Hilden, Germany) and XCI was determined by enzymatic methylation analysis of the human androgen receptor gene (AR, MIM313700) using the methylation-sensitive restriction enzyme HhaI and primers as described previously,[12] with minor methodological modifications. In brief, within the first exon, the *AR* gene contains a highly polymorphic short tandem repeat ($(CAG)_n$) in close proximity to several methylation-sensitive restriction sites. This allows the discrimination between both X-chromosome alleles by the CAG repeat sizes and the determination of their respective methylation status. In parallel to the methylation-sensitive digestion with HhaI, DNA of the same iPSC sample was treated identically to the digested sample, but no HhaI enzyme was added. This procedure of parallel treatment of the template DNA (HhaI digested and non-digested) allowed comparison and exact calculation of the degree of X-chromosome inactivation ratios.

Both, digested and non-digested DNA were amplified and fragment analysis was done by capillary electrophoresis on an ABI 3130xl Genetic Analyzer (Thermo Fisher Scientific, Waltham, MA, USA). Fragment sizes and peak area sizes were analysed using the GeneMapper™ software 6 (Thermo Fisher Scientific, Waltham, MA, USA). The XCI pattern was determined by calculating the ratios of the areas under the smaller ($XC_1$) and larger ($XC_2$) peaks as described previously:[13]

$$XCI\ (C_1) = \frac{\frac{XC_1^{digested}}{XC_1^{nondigested}}}{\frac{XC_1^{digested}}{XC_1^{nondigested}} + \frac{XC_2^{digested}}{XC_2^{nondigested}}} \quad (3)$$

$$XCI\ (C_2) = \frac{\frac{XC_2^{digested}}{XC_2^{nondigested}}}{\frac{XC_1^{digested}}{XC_1^{nondigested}} + \frac{XC_2^{digested}}{XC_2^{nondigested}}} \quad (4)$$

### Supplementary Tables

#### Supplementary Table 1: Antibodies for immunocytochemistry and immunohistochemistry

| **Marker** | **Source** | **Identifier** |
|---|---|---|
| Chicken polyclonal anti-beta III Tubulin | Abcam | Cat#ab41489; |
| Mouse monoclonal anti-peripherin- Alexa Fluor® 488 | Santa Cruz Biotechnology | Cat#sc-377093 |
| Mouse monoclonal anti-OCT3/4 | Santa Cruz Biotechnology | Cat#sc5279 |
| Mouse monoclonal anti-SSEA4 (clone MC813) | Cell Signaling Technologies | Cat#4755S |
| Mouse monoclonal anti-TRA-1-60 (clone TRA-1-60) | Millipore | Cat#901301 |
| Rabbit polyclonal anti-PAX6 | BioLegend | Cat#901301 |
| Mouse monoclonal anti-SOX2 (clone 245610) | R&D Systems | Cat#MAB2018 |
| Mouse monoclonal anti-FOXA2 | Santa Cruz Biotechnology | Cat#sc374376 |
| Rabbit polyclonal anti-SM22A | Abcam | Cat#ab14106 |

| | | |
|---|---|---|
| Human monoclonal anti-TRA-1-60-PE (clone REA157) | Miltenyi Biotec | Cat#130-100-347 |
| Human monoclonal anti-SSEA4-APC (clone REA101) | Miltenyi Biotec | Cat#130-098-347 |
| Rabbit monoclonal anti-LAMP1 (clone D2D11) | Cell Signaling Technologies | Cat#9091; RRID |
| Rabbit polyclonal anti-NaV1.8 | Alomone Labs | Cat#ASC-016 |
| Rat monoclonal anti-Substance P | Santa Cruz Biotechnology | Cat#sc-21715 |
| Goat polyclonal anti-TRPV1 | Santa Cruz Biotechnology | Cat#sc-12503 |
| Mouse monoclonal anti-PGP 9.5 (clone 31A3) | Bio-Rad | Cat#7863-1004 |
| Mouse monoclonal anti-Tom20 | Santa Cruz Biotechnology | Cat#G1819 |
| Rabbit-anti-peripherin | Merck Millipore | Cat#AB1530 |
| Donkey-anti-rabbit Cy3 AffiniPure | Jackson ImmunoResearch | Cat#711-165-152 |
| Goat anti mouse Alexa Fluor® 647-AffiniPure | Abcam | Cat#ab150115 |
| Mouse monoclonal anti-vimentin | Abcam | Cat#8978 |

**Abbreviations:** Cy3 = Cyanine 3, FOXA2 = Forkhead box protein A2, LAMP1 = against lysosomal-associated membrane protein 1, NaV1.8 = Voltage gated sodium channel 1.8, OCT3/4 = Octamer-binding transcription factor 3/4, PAX6 = Paired box 6, PGP9.5 = Protein gene product 9.5, SM22A = Smooth muscle protein 22-alpha, SOX2 = SRY-box transcription factor 2, SSEA4 = Stage-specific embryonic antigen-4, TRPV1 = Transient receptor potential vanilloid type 1.

#### Supplementary Table 2: List of TaqMan probes used

| **Marker** | **Source** | **Assay-ID** |
|---|---|---|
| GAPDH | Thermo Fisher Scientific | Hs02786624_g1 |
| TUBB3 (TUJ1) | Thermo Fisher Scientific | Hs00801390 _s1 |
| PRPH | Thermo Fisher Scientific | Hs00196608_m1 |
| POU4F1 (BRN3A) | Thermo Fisher Scientific | Hs00366711_m1 |
| NTRK1 (TRKA) | Thermo Fisher Scientific | Hs00176787_m1 |
| ISL1 | Thermo Fisher Scientific | Hs00158126_m1 |
| TRPV1 | Thermo Fisher Scientific | Hs00218912_m1 |
| TAC1 | Thermo Fisher Scientific | Hs00243225_m1 |
| SCN9A | Thermo Fisher Scientific | Hs00161567_m1 |
| SCN10A | Thermo Fisher Scientific | Hs01045137_m1 |

**Abbreviations:** GAPDH = glyceraldehyde-3-phosphate dehydrogenase, ISL1 = islet-1, NTRK1/TRKA = neurotrophic receptor tyrosine kinase 1, POU4F1/BRN3A = brain-specific homeobox/POU domain protein 3A, PRPH = peripherin, SCN9A = sodium voltage-Gated Channel Alpha Subunit 9, SCN10A = sodium voltage-gated channel alpha subunit 10, TAC1 = substance P, TRPV1 = transient receptor potential vanilloid type 1, TUBB3/TUJ1 = class III beta-tubulin.

#### Supplementary Table 3: Clinical characteristics of patient cohort

| | FD-1 | FD-2 | FD-3 |
|---|---|---|---|
| **Age, sex** | 28, m | 18, m | 25, f |
| **Genotype** | c.1096C>T hemizygous | c.568delG hemizygous | c.708G>C heterozygous |
| **Mutation type** | Nonsense | Frameshift mutation leading to premature stop codon, i.e. nonsense | Missense |
| **Cardiomyopathy** | Yes | No | No |
| **Nephropathy** | No | No | No |
| **FD-associated pain** | Attacks | No pain | Attacks |
| **FD- treatment** | Agalsidase-β | None | None |
| **GLA activity** | 0.5 (ref.: 3.4 – 13.0 nmol/h/ml) | 0.04 (ref.: 0.4 – 1.0 nmol/min/mg/protein) | 0.22 (ref.: 0.4 – 1.2 ng/ml/protein) |
| **Lyso-Gb3** | 57.7 (ref.: < 0.9 ng/ml) | 241 (ref.: < 20.1 ng/ml) | 11.5 (ref.: < 0.9 ng/ml) |

**Abbreviations:** GLA = alpha-galactosidase A, f = female, FD-1, 2, 3 = patients with Fabry disease, lyso-Gb3 = lyso globotriaosylceramide, m = male.

### Supplementary Figures

#### Supplementary Figure 1

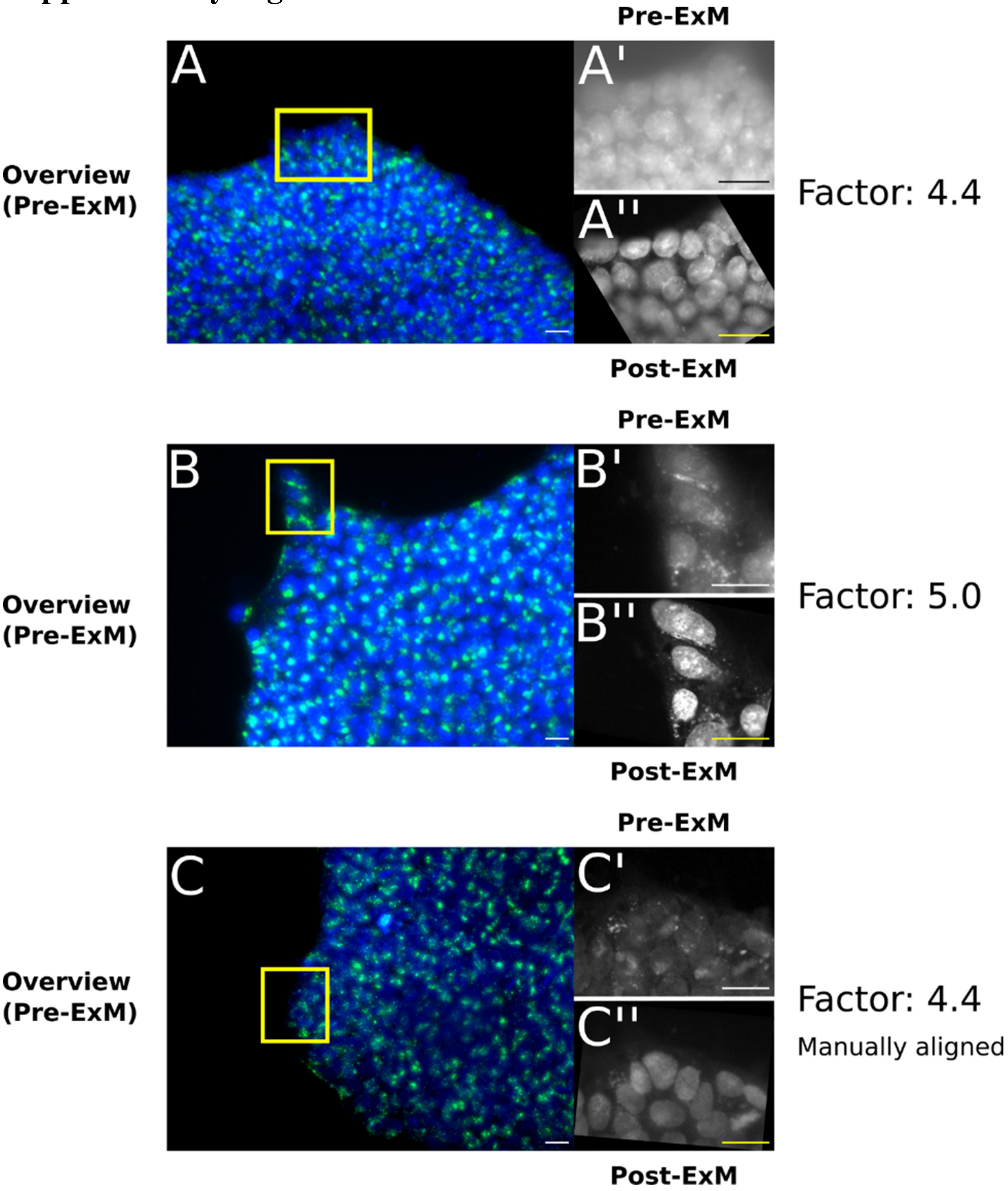

#### Supplementary Figure 2

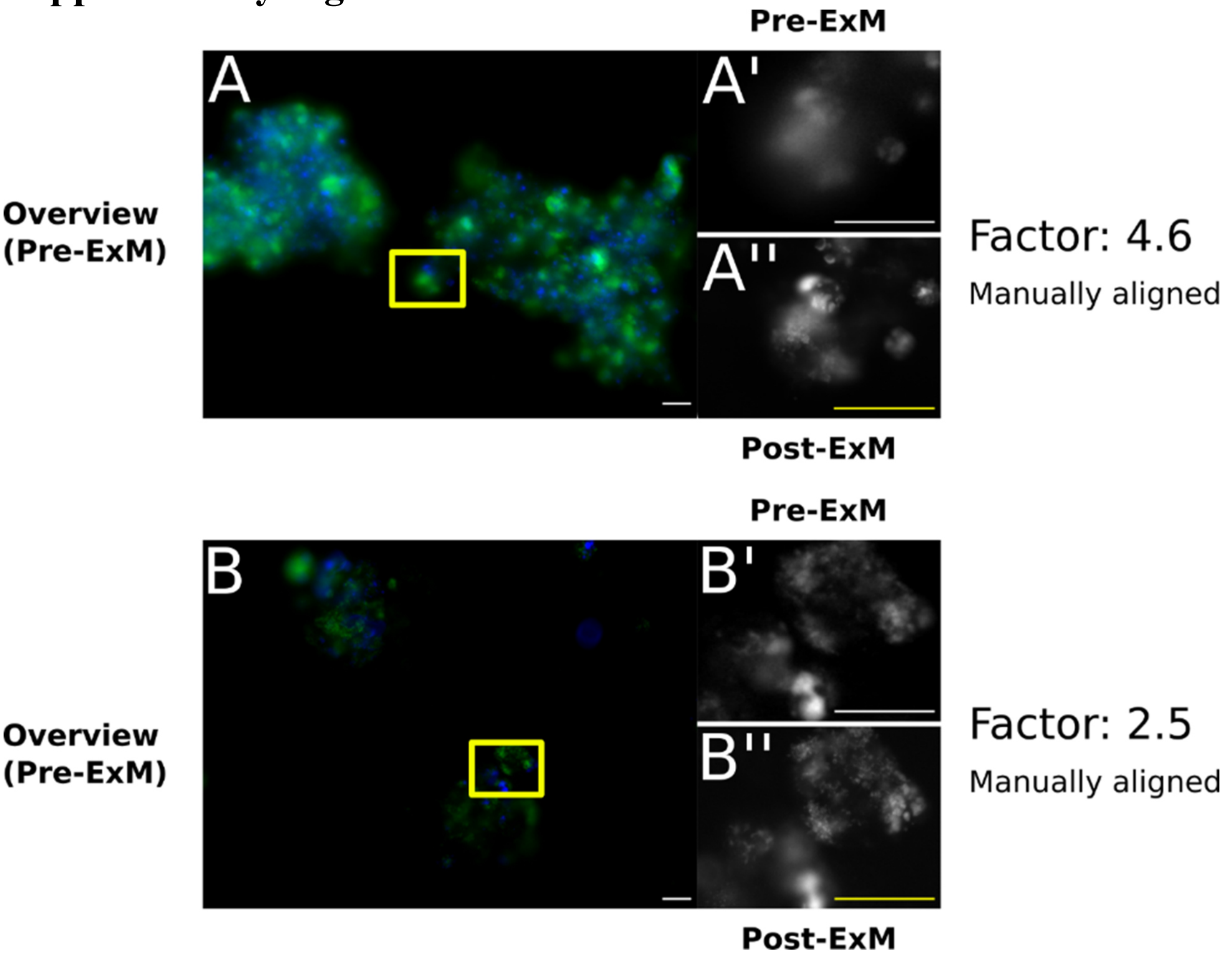

### Supplementary Figure 3

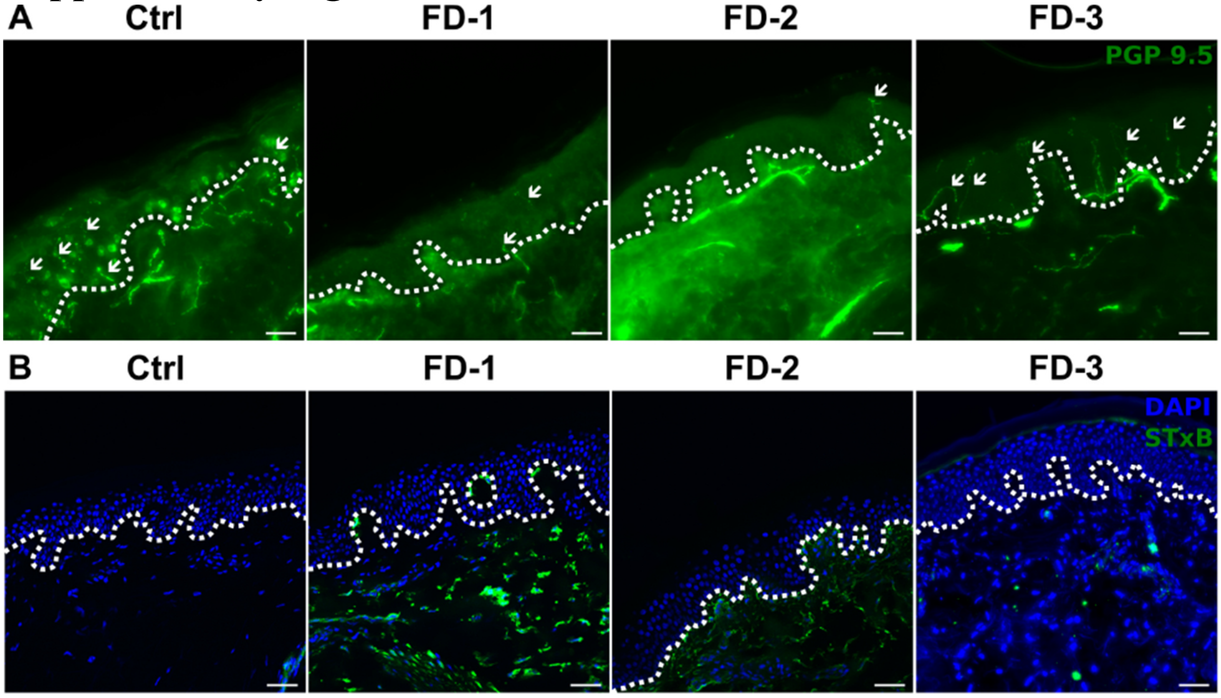

#### Supplementary Figure 4

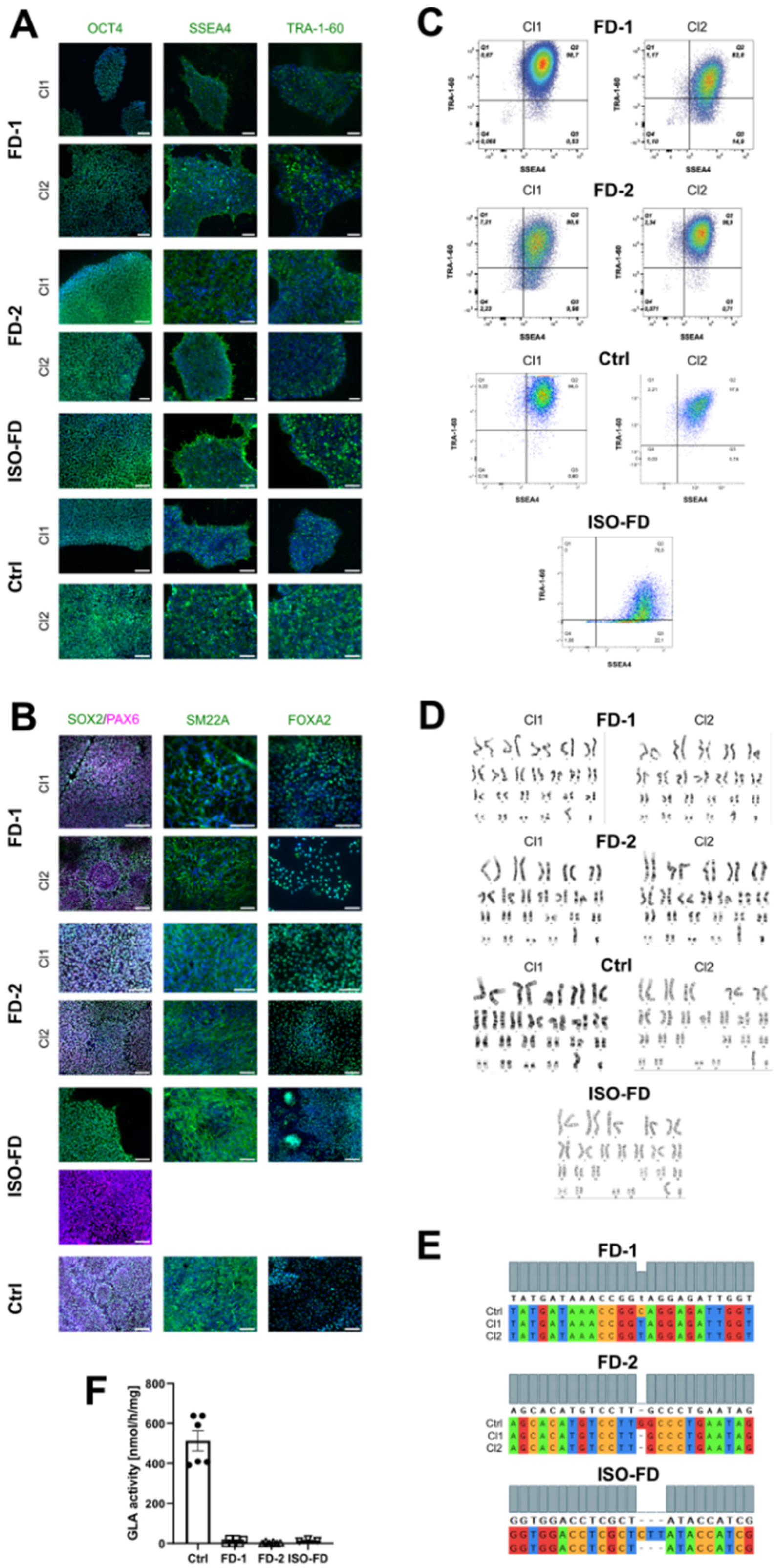

### Supplementary Figure 5

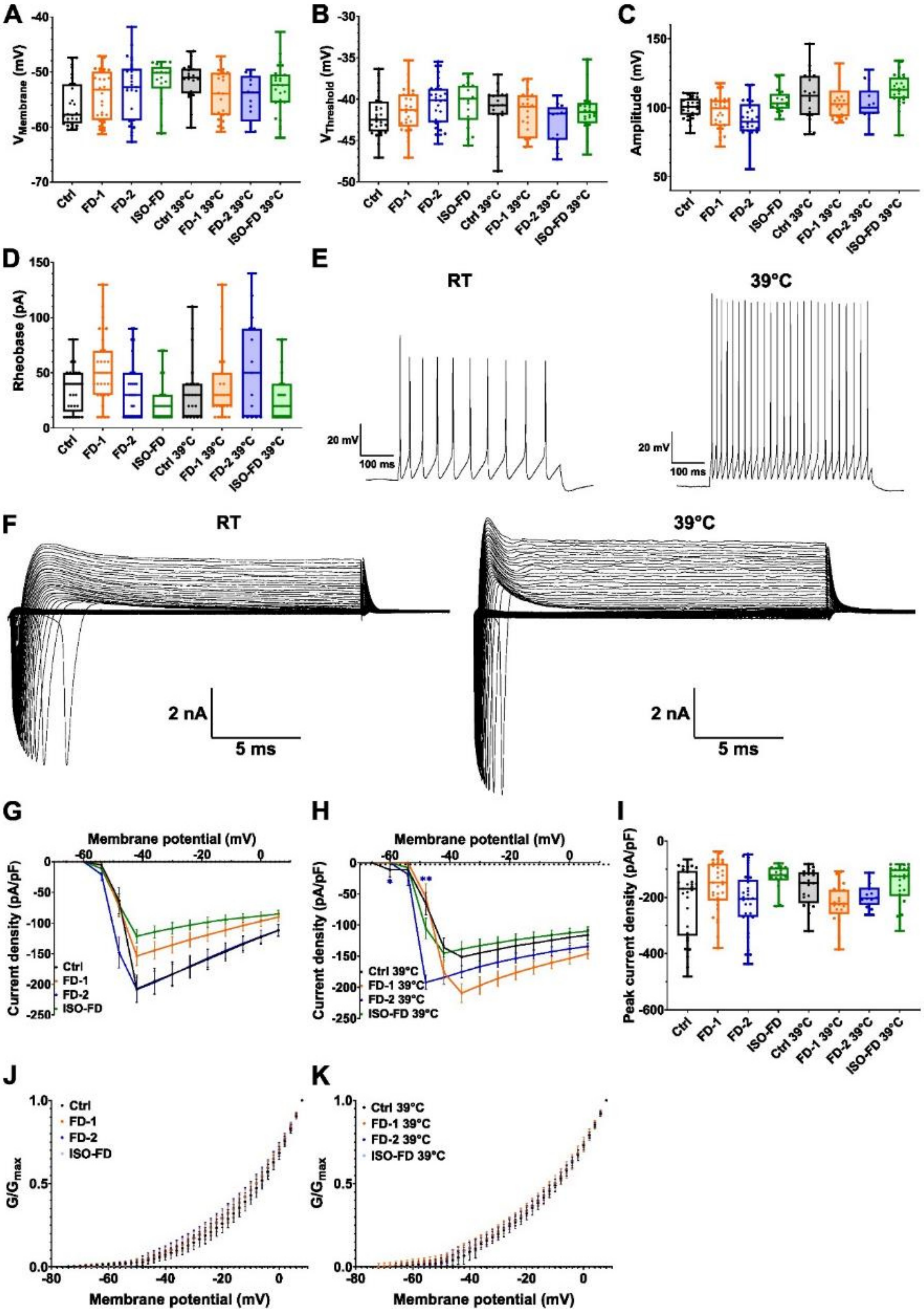

### Legends Supplementary Figures

#### Supplementary Figure 1: Expansion microscopy in iPSC

**(A – C)** Photomicrographs of Ctrl, FD-1, and FD-2 iPSC before expansion using DAPI (blue) and LAMP1 (green) as nuclear and lysosomal markers. The yellow rectangle indicates the region of interest (ROI) used for correlation after expansion. Scale bars: 25 µm. **(A' – C')** ROI used for calculation of expansion factor. Scale bars: 25 µm. **(A'' – C'')** ROI after expansion with the expansion corrected scale. Scale bars: 25 µm. All images: Min/Max values were adjusted for each channel for visualization. For Ctrl and FD-1, expansion factor was calculated automatically, whereas for FD-2, expansion factor was calculated manually, due to technical issues. Acquired with Leica Dmi8 (widefield). **Abbreviations:** ExM = Expansion microscopy, Ctrl = control, DAPI = 4',6-diamidino-2-phenylindole, FD-1, 2 = Fabry patients, iPSC = induced pluripotent stem cells, LAMP1 = lysosomal-associated membrane protein 1, ROI = region of interest.

#### Supplementary Figure 2: Expansion microscopy in neurons

**(A, B)** Photomicrographs of FD-1 and FD-2 iPSC-derived sensory neurons before expansion using DAPI (blue) and LAMP1 (green) as nuclear and lysosomal markers. The yellow rectangle indicates the ROI used for correlation after expansion. Scale bars: 25 µm. **(A'**, **B')** ROI used for calculation of expansion factor. Scale bars: 25 µm. **(A'', B'')** ROI after expansion with the expansion corrected scale. Scale bars: 25 µm. All images: Min/Max values were adjusted for each channel for visualization. For FD-1 and FD-2, expansion factor was calculated manually, due to technical issues with the python script. Acquired with Leica Dmi8 (widefield). **(**B), (B'),

(B'') For FD-2, expansion factor was calculated with expanded images and images of the ROI after shrinking the gel back in PBS overnight, since the original pre-expansion site could not be identified. However, this lead to a discrepancy of expansion factor with only 2.5x. Incomplete shrinking of the gel accounted for this artifact. Since the same expansion protocol for iPSC and neurons was applied and FD-1 and FD-2 neurons were expanded within one batch, we estimate a total expansion factor in the range of 4.6x as shown for FD-1. **Abbreviations:** ExM = expansion microscopy, Ctrl = control, DAPI = 4',6-diamidino-2-phenylindole, FD-1, 2 = Fabry patients, iPSC = induced pluripotent stem cells, LAMP1 = lysosomal-associated membrane protein 1, ROI = region of interest.

#### Supplementary Figure 3: Assessment of skin sections obtained from Fabry patients and healthy control

(**A**) Intraepidermal nerve fibre density was reduced in FD-1 and FD-2, but was in the normal range in FD-3 and Ctrl. White arrows point at individual nerve fibres. Scale bars: 50 µm. (**B**) Dense dermal Gb3 accumulations (green) were found in skin samples of all three patients while absent in Ctrl. Scale bars: 50 µm. The white dotted line marks epidermis-dermis border in A and B. We used the following instruments and post-processing steps for the generation of our images: In (A), images were acquired using widefield Z-stacks and maximum intensity projections. Exposure time was identical for all channels and images. Acquired with Axio Imager.M2 (widefield). (B): Images were acquired using Z-stacks and maximum intensity projections with an Apotome.2 device. Exposure time for STxB::555 in the Ctrl sample was higher than in FD-1 and FD-2. Min/Max pixel values were adjusted in an identical way.

Acquired with Axio Imager.M2 and Apotome.2. **Abbreviations:** Ctrl = control, FD-1, 2 = Fabry patients, Gb3 = Globotriaosylceramide, STxB = Shiga toxin 1

#### Supplementary Figure 4: Characterization of iPSC and isogenic control

**(A)** Expression of pluripotency markers OCT4, SSEA4, and TRA-1-60 was confirmed in both clones of FD-1, FD-2, Ctrl, and for the ISO-FD clone. Scale bars: 100 µm. Acquired with Axiophot (widefield; FD-1), and Axio Imager.M2 (widefield; FD-2 and Ctrl). **(B)** FD-1, FD-2, ISO-FD, and Ctrl clones were differentiated into ectodermal, mesodermal, and endodermal cells to further investigate pluripotency. All clones expressed the respective germ layer markers SOX2/PAX6 (ectoderm), SM22A (mesoderm), and FOXA2 (endoderm). Scale bars: 50 µm. Acquired with Axiophot (widefield; FD-1), and Axio Imager.M2 (widefield; FD-2, ISO-FD, Ctrl). (A) + (B): Min/Max pixel values were adjusted for visualization for each channel. Nuclei were visualized using DAPI. **(C)** Quantitative expression of pluripotency markers was confirmed via FACS for Ctrl, FD-1, FD-2, and ISO-FD lines. **(D)** Genomic integrity was confirmed via G-banding for FD-1, FD-2, ISO-FD, and Ctrl lines. **(E)** Disease specific mutations were analysed in FD-1, FD-2, and ISO-FD lines. Ctrl-iPSC were used as reference. **(F)** GLA activity was reduced in FD-1, FD-2 iPSC (each n = 2 clones), and ISO-FD (n = 1 clone) compared to Ctrl-iPSC (n = 1 clone). Data are represented as mean ± SD. **Abbreviations:** Ctrl = control, DAPI = 4',6-diamidino-2-phenylindole, FD-1, FD-2 = patients with Fabry disease, FOXA2 = forkhead box protein A2, GLA = alpha-galactosidase A, iPSC = induced pluripotent stem cells, ISO-FD = isogenic Fabry line, OCT4 = octamer-binding transcription factor 4, PAX6 = paired box 6,

SM22A = smooth muscle protein 22-alpha, SOX2 = SRY-box transcription factor 2, SSEA4 = stage-specific embryonic antigen-4.

#### Supplementary Figure 5: Electrophysiology

(**A-D**) No differences in Vmembrane (**A**), Vthreshold (**B**), action potential amplitudes (**C**), and rheobase currents (**D**) between Ctrl, FD-1, FD-2 and ISO-FD neurons at RT and 39°C. Data are represented as box-and-whisker plots with dots as individual values. Box width indicates the first and third quartiles, horizontal line indicates the median, and whiskers indicate the lowest and highest values. One-way ANOVA followed by Sidak's multiple comparison correction. (**E**) Representative current-clamp recordings at RT and 39°C. **(F**) Exemplary voltage-clamp recordings of inward voltage-gated sodium currents and outward voltage-gated potassium currents at RT and 39°C. (**G**) No differences in voltage-gated sodium current densities between Ctrl, FD-1, FD-2 and ISO-FD neurons at RT. Data are represented as mean ± SEM. Two-way ANOVA followed by Sidak's multiple comparison correction. (**H**) At 39°C, voltage-gated sodium current densities are increased in FD-2 neurons at -60 mV and -48 mV compared to Ctrl neurons. Data are represented as mean ± SEM. Two-way ANOVA followed by Sidak's multiple comparison correction. (**I**) No differences in peak current densities between Ctrl, FD-1, FD-2, and ISO-FD neurons at RT and 39°C. Data are represented as box-and-whisker plots with dots as individual values. Box width indicates the first and third quartiles, horizontal line indicates the median, and whiskers indicate the lowest and highest values. Kruskal-Wallis test followed by Dunn's multiple comparison correction. (**J**) Steady-state activation curves of voltage-gated potassium channels were comparable between the cell lines at RT ($V_{1/2}$: Ctrl: -8.94 mV; FD-1: -

10.21 mV; FD-2: -10.29 mV; ISO-FD: -10.52 mV). Data are represented as mean ± SD. (**K**) Steady-state activation curves of voltage-gated potassium channels were comparable between the cell lines at 39°C ($V_{1/2}$: Ctrl: -11.66 mV; FD-1: -12.25 mV; FD-2: -12.32 mV; ISO-FD: -11.71 mV). Data are represented as mean ± SD. For (A-D): For Ctrl (n = 2 clones; RT: clone 1 = 8 cells, clone 2 = 21 cells; 39°C: clone 1 = 16 cells, clone 2 = 4 cells), FD-1 (n = 2 clones; RT: clone 1 = 15 cells, clone 2 = 16 cells; 39°C: clone 1 = 18 cells, clone 2 = 5 cells), FD-2 (n = 2 clones; RT: clone 1 = 11 cells, clone 2 = 18 cells; 39°C: clone 1 = 15 cells), and ISO-FD (n = 1 clone; RT: 19 cells, 39°C: 26 cells) pooled data obtained from ≥ 3 individual differentiations were used. For (E-G): For Ctrl (n = 2 clones; RT: clone 1 = 8 cells, clone 2 = 21 cells; 39°C: clone 1 = 16 cells, clone 2 = 4 cells), FD-1 (n = 2 clones; RT: clone 1 = 15 cells, clone 2 = 16 cells; 39°C: clone 1 = 16 cells, clone 2 = 5 cells), FD-2 (n = 2 clones; RT: clone 1 = 11 cells, clone 2 = 18 cells; 39°C: clone 1 = 14 cells), and ISO-FD (n = 1 clone; RT: 16 cells, 39°C: 26 cells) pooled data obtained from ≥ 3 individual differentiations were used. **Abbreviations:** Ctrl = control; FD-1, 2 = Fabry patients; RT = room temperature; VMembrane = membrane potential; Vthreshold = threshold potential. *$p < 0.05$; **$p < 0.01$.

### Legends Supplementary Videos

#### Supplementary Video 1: Neuronal firing induced by KCl incubation

Neuronal activity was induced by application of KCl (right video) compared to non-treated neurons (left video). Normalized intensity was plotted against the acquisition time and an increase of the calcium indicator dye Fluo 8-AM can be seen in the KCl treated group after adding at 60 seconds. Scale bars: 25 µm. **Abbreviations:** KCl = potassium chloride.

#### Supplementary Video 2: Mitochondrial traffic jam

Time lapse video of mitochondrial tracking (MitoTracker, purple) after incubation with sphinganine (green) of FD-1 neurons hinting to impairment of mitochondrial mobility due to sphinganine accumulations. Scale bar: 25 µm.

#### Supplementary Video 3: Mitochondrial traffic jam – detailed view "G"

Detailed view of the region of interest "G" from Supplementary Video 2 showing that mitochondria (purple) and sphinganine (green) interact, which may interfere with normal mitochondrial mobility. Scale bar: 5 µm.

#### Supplementary Video 4: Mitochondrial traffic jam – detailed view "H"

Detailed view of the region of interest "H" from Supplementary Video 2 showing that mitochondria (purple) and sphinganine (green) interact possibly hindering mitochondrial mobility. Scale bar: 5 µm.
